# De-novo promoters emerge more readily from random DNA than from genomic DNA

**DOI:** 10.1101/2025.08.25.672121

**Authors:** Timothy Fuqua, Andreas Wagner

## Abstract

Promoters are DNA sequences that help to initiate transcription. Point mutations can create de-novo promoters, which can consequently transcribe inactive genes or create novel transcripts. We know little about how de-novo promoters emerge in genomic DNA, especially compared to random DNA that has never been subjected to selection. Here, we assayed the promoter activity of 17,129 random, synthetic DNA sequences and 91,866 *E. coli* genomic DNA sequences. Genomic DNA encodes ~1.3 times more promoters than random DNA. We then studied 584,573 point mutations in 225 random and 60 genomic sequences, and asked how they cause the emergence of de-novo promoters. We find that de-novo promoters emerge ~3 times more readily from random DNA than from genomic DNA. The reason is that the genome contains fewer proto-binding sites for transcriptional activators than random DNA. Our work shows that the evolutionary history of a DNA sequence introduces substantial biases in its evolutionary potential, especially in the likelihood that mutations create new and potentially adaptive transcripts.

## Introduction

Gene regulation enables organisms to respond to their environment^1^. It is controlled on the level of transcription by promoter sequences on DNA^2^. In prokaryotes, canonical promoters encode binding sites for sigma (σ) factors that help RNA polymerase bind near a gene^3–6^. Promoters may also encode binding sites for transcription factors (TFs) to activate or repress transcription, depending on the environment^1,7^. Mutations in promoters can have profound impacts on their activity^7–12^. In addition, mutations can cause the emergence of new promoters, which can lead to evolutionary innovations, either by changing the regulation of existing genes^13–15^, or by creating de-novo transcripts^16,17^.

Promoters are simple DNA sequences. For example, ~10% of randomly synthesized DNA sequences have promoter activity, depending on length and AT-content^18–21^. Furthermore, promoters can emerge from non-promoter DNA sequences by single nucleotide mutations^18,19^. Such mutations can cause a gain of binding sites for a σ factor or for a TF that activates transcription^18,22–24^. They can also cause a loss of a binding site for a TF that represses transcription, or a loss of a transcriptional terminator sequence^7,23,24^.

Unlike random synthetic DNA sequences, genomes are products of Darwinian evolution that encode various functions^25,26^. For example, natural selection has helped to create the promoters that are necessary to transcribe protein coding genes. However, selection also acts against promoters that coincidentally exist inside of coding DNA^27–29^. Computational analyses reveal that the *E. coli* genome overall contains fewer promoter-like sequences than random DNA^18,19,30^, and its protein-coding DNA is impoverished in codons that resemble parts of σ-factor binding sites^11,30^. Furthermore, regulatory proteins also help to reduce transcription that starts inside coding DNA through repression and premature transcriptional termination^31–33^. Thus, there are simultaneous selective pressures to preserve the promoters of thousands of functional genes^34^ and to remove promoters inside these genes^31–33^. This dichotomy may influence how new promoters emerge in the *E. coli* genome. We quantify this influence by comparing how likely it is that promoters emerge *de novo* in random DNA and genomic DNA.

Specifically, we experimentally explore the propensity of both random and genomic DNA to be and become promoters. We find that genomic DNA is approximately ~1.3 times more likely to encode promoter activity than random DNA, regardless of whether it originates from inside or outside coding DNA. However, upon mutation, promoters emerge ~3 times more readily in random DNA than in genomic DNA. We attribute these differences to two general observations. First, the genome encodes almost ~4 times fewer cryptic binding sites for σ factors and TFs, which lowers the likelihood that mutations create new such sites. Second, genomic DNA is de-repressed relative to random DNA. This de-repression may explain why there are more promoters in protein-coding genes than in random DNA. Overall, our study reveals how the different sequence backgrounds of functional genomic DNA and random DNA can bias the emergence of promoters, which by extension, may influence the emergence of evolutionary adaptations and innovations.

## Results

### Random DNA is very likely to encode a promoter, and genomic DNA even more so

We asked how likely it is that a random DNA sequence encodes a bacterial promoter, and how that likelihood compares to that of a genomic DNA sequence. To this end, we first created a *random DNA library* by synthesizing a pool of oligonucleotides (IDT, USA), each 150bps in length, where each position has a 25% chance of encoding an A, T, C, or G (**Fig 1A**). Second, we created a *genomic DNA library* by fragmenting the *E. coli* genome into 100-300 bp pieces using ultrasonication (**Methods**). We cloned both of these libraries into a dual-reporter plasmid^35^ (pMR1) between a green (red) fluorescent protein coding sequence downstream (upstream) of a library sequence, and transformed *E. coli* cells with the resulting plasmid library. Transformed bacteria express green fluorescent protein (GFP) or red fluorescent protein (RFP) when a library insert harbors a promoter on the top or bottom DNA strand, respectively. To map the fluorescence of each bacterium to its respective plasmid insert, we used Sort-Seq^8,23,24,36^ to separate bacteria with a cell sorter (BD Biosciences, FACSAria III) into eight fluorescence bins corresponding to none, weak, moderate, and strong GFP or RFP expression (**Fig 1B**). We then bulk-sequenced the plasmid’s DNA inserts from each bin, and calculated fluorescence scores in arbitrary units (a.u.) for each insert based on read counts. These scores range from 1.0 a.u. (no expression) to 4.0 a.u. (highest expression, **Methods**).

**Figure 1.**
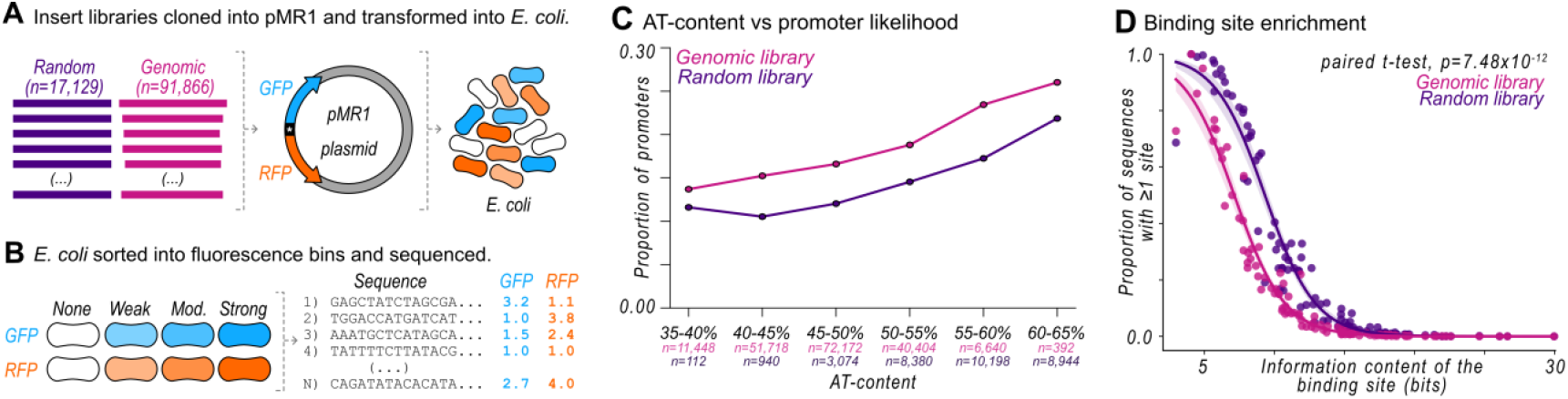
The genome is enriched with active promoters relative to random DNA. **(A)** We cloned the random library of 150 bp N-mer sequences (n=17,129, purple), and the genomic library of 100-300 bp sequences (n=91,866, magenta) into the dual-reporter plasmid MR1 (pMR1), which drives the expression of green fluorescent protein (GFP, teal) from inserts on the top DNA strand, and that of red fluorescent protein (RFP, orange) on the bottom strand. We transformed *E. coli* cells with the plasmid libraries. **(B)** We sorted the bacterial libraries into fluorescence bins at four fluorescence strengths: none, weak, moderate, and strong for both GFP and RFP (eight bins total) with a cell-sorter. We bulk-sequenced the library inserts from each bin and calculated fluorescence scores in arbitrary units (a.u.) ranging between one (none) and four (strongest) (**Methods**). **(C)** The probability that a DNA sequence in the random (purple) and genomic (magenta) libraries is a promoter relative to its AT-content. **(D)** For 102 position-weight matrices (PWMs) for transcription factors and sigma (σ) factors, we plot the percentage of sequences in each library (purple: random, magenta: genome) that encode at least one putative factor binding site (vertical axis) against the respective PWM’s information content in bits. We test for equality of the frequency distributions between the random and genome libraries with a paired t-test (p=7.48×10^−12^). See **Source Data**.

We screened 17,129 unique 150bp sequences from the random library, with an average AT-content of ~57±6%. We defined a sequence as a promoter if it drove fluorescence expression of at least 1.5 arbitrary units (a.u.). At this threshold ~13% of sequences with an AT-content of 45–55% function as promoters on either strand (**Fig 1C**), a percentage that aligns with previous studies reporting ~5%^20^, ~10%^18^, and ~10-20%^19^ of sequences in similar random DNA libraries encoding promoters on a single strand. Overall, 33 percent (5,694) of the 17,129 random sequences encode a promoter, with 27 percent (4,649) driving expression from the top strand, 4 percent (684) from the bottom strand, and 2 percent (361) from both strands (**Fig S1E**).

From the genomic library, we screened 91,866 unique sequences that collectively cover the genome ~2.7× fold ^37^ (**Fig S1B**). These sequences are ~130±23 bp in length (**Fig S1A**), with an average AT-content of ~47±5%. We note that these genomic sequences were integrated into the expression plasmid in either genetic orientation, such that a sequence from the top strand of the genome may be located on the top or bottom strand of the plasmid, and vice-versa (**Methods** and **Fig S9**). Overall, 32 percent (29,116) of the 91,866 genomic sequences encode a promoter. More specifically, 25 percent (23,012) of sequences drive expression from the top strand, 5 percent (4,394) from the bottom strand, and 2 percent (1,710) from both strands (**Fig S1E**).

The likelihood that DNA encodes a promoter increases with AT-content^20^, which differs between our two libraries (**Fig S1F**). To control for AT-content when comparing the incidence of promoters, we binned all random and genomic sequences into one of 6 intervals spanning 5% AT-content, and calculated the proportion of promoters for sequences in each bin. For both libraries, the proportion of promoters increases with AT-content (**Fig 1C**), but it is greater in genomic sequences for all bins. Overall, genomic sequences are ~1.3 times more likely to encode a promoter than random sequences (average ~19.0% vs ~14.6%).

Most known genomic promoters are products of natural selection to transcribe DNA, but selection also acts to remove promoters inside of protein-coding genes^11,18,19,30–33^. One might thus expect that intergenic DNA is more likely and intragenic DNA less likely to encode a promoter than expected by chance alone, i.e., in random DNA. To find out, we isolated 964 intergenic DNA sequences mapping exclusively to non-coding DNA, and 80,622 intragenic DNA sequences mapping exclusively to coding DNA from the genomic library. Unexpectedly, we found that DNA originating from *both* non-coding or coding DNA is more likely to encode promoters than random DNA (**Fig S2A,B**). The difference is greater for non-coding DNA (~25% vs ~15%). This is expected, because most known promoters occur in non-coding DNA (see **Table S1** for matched promoters from the RegulonDB database^38^). However, we had not expected that sequences exclusively inside coding DNA are also more likely to encode a promoter than random sequences (~19% vs ~15%). Many of these promoters are on the antisense strands of these genes (**Fig S2C**), which is consistent with previous reports^11,39^.

To exclude the possibility that the observed differences can be explained by differences in sequence length, we subsampled genomic sequences that are exactly 150bps long, just like the sequences in the random library. These 150bp genomic sequences are ~1.5 times (23% vs 15%, similar to ~1.3 times) more likely to encode a promoter than random sequences (**Fig S1E** and **Fig S2D**). Additionally, the correlation between sequence length and expression level in the genomic library is very small (Pearson’s r=0.01, p=4.91×10^−6^, **Fig S1C**), suggesting that length is not biasing these results. See **Fig S2** for additional analyses.

Altogether these analyses demonstrate that the genome’s enrichment of promoters does not just stem from canonical intergenic promoters, but also from intragenic sequences, particularly those driving antisense transcription.

### The genomic library contains fewer predicted activating and repressing sites than the random library

We asked whether enrichment of particular transcription factor (TF) and sigma (σ) factor binding sites in the libraries could explain why genomic DNA is more likely to encode a promoter than random DNA. To this end, we obtained a non-exhaustive set of 102 position weight matrices (PWMs) for 93 TFs and 9 σ factors (see **Methods**). A PWM is a matrix derived from protein-DNA binding experiments that represents the frequency of each nucleotide at every position within a TF’s binding site. A PWM has an information content, which reflects the spectrum of DNA sequences that a TF can bind. If a PWM has high information content, the corresponding TF binds few sequences and is thus highly specific. In addition, a PWM can be used to derive a predicted protein binding strength from a query sequence. If this strength is greater than a particular threshold, we call the query sequence a *predicted binding site* for a TF or σ factor (**Methods**).

For each PWM, we determined the percentage of library sequences that contain at least one predicted binding site. Not surprisingly, binding sites with low information content are more frequent in random DNA (**Fig 1D**). This is because such sites are expected to occur more often in random DNA by chance alone. While binding site frequency also varies with information content in genomic DNA (**Fig 1D**), binding site frequencies are on average ~10 percent lower in genomic DNA (~16% vs ~26%, paired t-test, p=7.48×10^−12^, see **Fig S1G** for details). TFs can either activate or repress transcription^40,41^ but σ factors exclusively activate transcription^23^. Together with our finding that genomic DNA is ~1.3 times more likely to encode a promoter, the relative scarcity of predicted sites in the genomic library implies that random DNA contains more binding sites that repress transcription (for brevity, it is more “repressed”) than genomic DNA, leading to less promoter activity overall.

### De-novo promoters emerge more readily from random DNA than from genomic DNA

To understand how promoters emerge from genomic DNA and how that compares to random DNA, we used fluorescence-activated cell sorting (FACS) to isolate 225 DNA sequences from the random library and 60 DNA sequences from the genomic library that drive neither GFP nor RFP expression (**Methods**). We call these sequences random and genomic *parent* sequences, respectively. For each of them, we created a mutagenesis library of mutant *daughter* sequences using an error-prone polymerase chain reaction (**Methods, Fig 2A**). We cloned each of the resulting 285 (60+225) mutagenesis libraries into the expression plasmid pMR1, and quantified the expression driven by each daughter sequence using Sort-Seq **(Methods)**. We normalized the fluorescence expression level of each daughter to the interval 1.0 a.u. and 4.0 a.u. (highest expression) (**Methods**).

**Figure 2.**
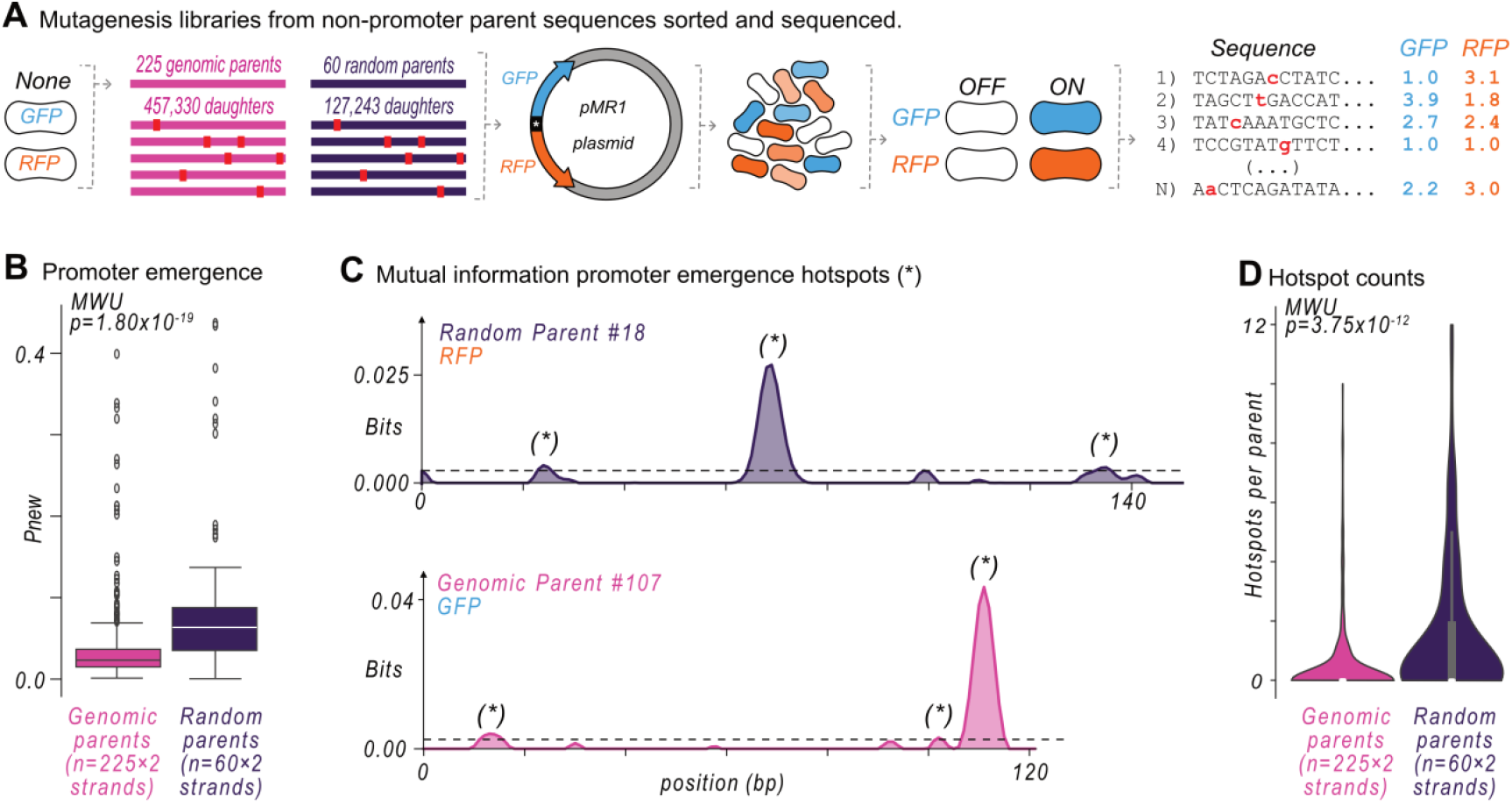
Promoters emerge more readily from random DNA than from genomic DNA. **(A)** We used fluorescence-activated cell sorting (FACS) to isolate sequences in the random and genomic libraries without promoter activity. From these *parent* sequences, we created mutagenesis libraries of *daughter* sequences, pooled the parents and their daughters, cloned them into the pMR1 plasmid, and transformed *E*.*coli* cells with the resulting plasmid libraries. We used Sort-Seq to partition the *E. coli* cells into four different fluorescence bins: GFP off, GFP on, RFP off, and RFP on, and sequenced the DNA inserts from each fluorescence bin to calculate sequence-specific fluorescence scores (see Methods). **(B)** For each parent and its respective daughters, we calculated *Pnew*, the proportion of daughters with a fluorescence score ≥ 1.5 arbitrary units (a.u.). We plotted *Pnew* for the random vs the genomic parents. *Pnew* is significantly greater for random sequences (two-tailed Mann-Whitney U [MWU] test, p=1.80×10^−19^). The center line shows the median, the box the interquartile range (IQR), and whiskers span ±1 standard deviation. See **Fig S4** for additional analyses on *Pnew* vs other sequence features (i.e. AT-content, number of mutations). **(C)** We calculated at each nucleotide position *i* along a sequence the mutual information *I*_*i*_ *(b,j)* between nucleotide identity (*j* = *A, T, C or G*), and fluorescence scores (*f* = *1*.*0 – 4*.*0 a*.*u*.) in information theoretical units (bits). The x-axis shows the position *i*, and the y-axis the sum of mutual information for each nucleotide *j* at this position. The dashed line indicates the 0.0025 bit threshold above which we call a mutual information peak a “hotspot” (asterisk). Top: the mutual information for *random* parent #18 and the propensity of mutations to create promoter activity in the bottom (RFP) strand of this parent. Bottom: mutual information for *genome* parent #107 and the propensity of mutations to create promoter activity in the top (GFP) strand. **(D)** The number of hotspots in each parent in the random mutagenesis library vs the genome mutagenesis library (two-tailed Mann Whitney U [MWU] test, p=3.75×10^−12^). See **Source Data**.

Overall, the *random mutagenesis libraries* contain a total of ~1.3×10^5^ mutant daughter sequences, i.e., on average 2,121 mutant daughters for each of the 60 parents. The *genomic mutagenesis libraries* contain ~4.6×10^5^ mutant daughter sequences, on average 2,033 mutant daughters for each of the 225 parents (see **Fig S3** for library details).

We first quantified the probability *Pnew* that a parent can acquire de-novo promoter activity by dividing, for each parent, the number of daughter sequences that drive gene expression (fluorescence ≥ 1.5 a.u) by the total number of daughters^23,24^. For example, *random parent #18* has 772 mutant daughters, of which 252 have promoter activity on the top (GFP) strand. Thus, *Pnew* for this parent equals ~0.33 (252 /772). We found that *Pnew* is ~3 times higher for random parents than for genomic parents (median *Pnew* =0.092 vs 0.029, two-tailed MWU test, p=1.80×10^−19^) (**Fig 2B**). This difference is significant regardless of the number of point mutations in the daughters (**Fig S4A**), number of daughter sequences (**Fig S4E**), and the length of the parent sequence (**Fig S4F**). Pnew does not significantly increase with AT-content for the random parents (Pearson correlation, p=0.241) but does weakly for the genomic parents (Pearson correlation, r=0.172, p=2.48×10^−4^) (**Fig S4C**). Nevertheless, random parents are still more likely to create de-novo promoters compared to genomic parents when grouped by their AT-contents (**Fig S4D**).

In sum, new promoters emerge ~3 times more readily from random DNA than from genomic DNA.

### Genomic DNA harbors fewer emergence hotspots than random DNA

In previous work, we had demonstrated that new promoters typically emerge from “hotspot” regions in a parent sequence^23,24^. To identify these hotspots in each parent, we calculated for each nucleotide position *i* along the parent sequence, the mutual information *(I*_*i*_*(b,j))* between nucleotide identity (*j* = `A,T,C,G`), and fluorescence expression (*f* = *1*.*0 – 4*.*0 a*.*u*.) (**Methods**)^8^. This calculation creates a profile of mutual information peaks and valleys for each parent sequence, depicting where promoters can – and cannot – emerge (**Fig 2C**)^7– 10,12,23,24^. We define a “hotspot” as a mutual information peak greater than 0.0025 bits (**Methods**). See **Fig S5** for mutual information plots for all parents with at least one hotspot.

Remarkably, the average genomic parent has ~3.8 times less hotspots than the average genomic parent (~0.4 hotspots vs ~1.5 hotspots; two-tailed MWU test, p=3.75×10^−12^ **Fig 2D**). Modifying our definitions of a hotspot does not change this conclusion (**Fig S6**). For each parent, the number of hotspots strongly correlates with the probability of promoter emergence *Pnew* (Pearson r=0.750 and 0.717, p=5.84×10^−23^ and 1.82×10^−72^ for random and genomic parent, respectively, **Fig S4B**). Random and genomic parents with the same number of hotspots have virtually the same value of *Pnew* (**Fig S4B**).

In sum, point mutations can create de novo promoters in both random and genomic DNA, but they are about three times more likely to do so in random DNA. The reason is that random DNA harbors many more unique hotspot locations than genomic DNA in which such promoters can emerge (**Fig 2)**.

### The *E. coli* genome is impoverished in proto-sites for activating factors

Mutual information hotspots occur where sequences either *gain* new binding sites for activating TFs and σ factors, or where they *lose* repressing TF sites^23,24^. We first focused on the former case using a non-exhaustive set of 9 σ factor and 93 TF binding sites (102 PWMs total). Specifically, for those daughters that gain a predicted site by mutation, we used a two-tailed Mann-Whitney U test to ask whether they also experienced a significant increase in gene expression of ≥ 0.1 a.u. (**Methods**). For example, *random parent #37* gains de-novo promoter activity when mutations create a binding site (−10 box) for the canonical housekeeping σ^70^ factor (**Fig 3A,B**) (MWU test, q=1.52×10^−26^). We call newly created sites associated with increasing transcriptional activity *activating sites*. See **Fig S7** for additional examples.

**Figure 3.**
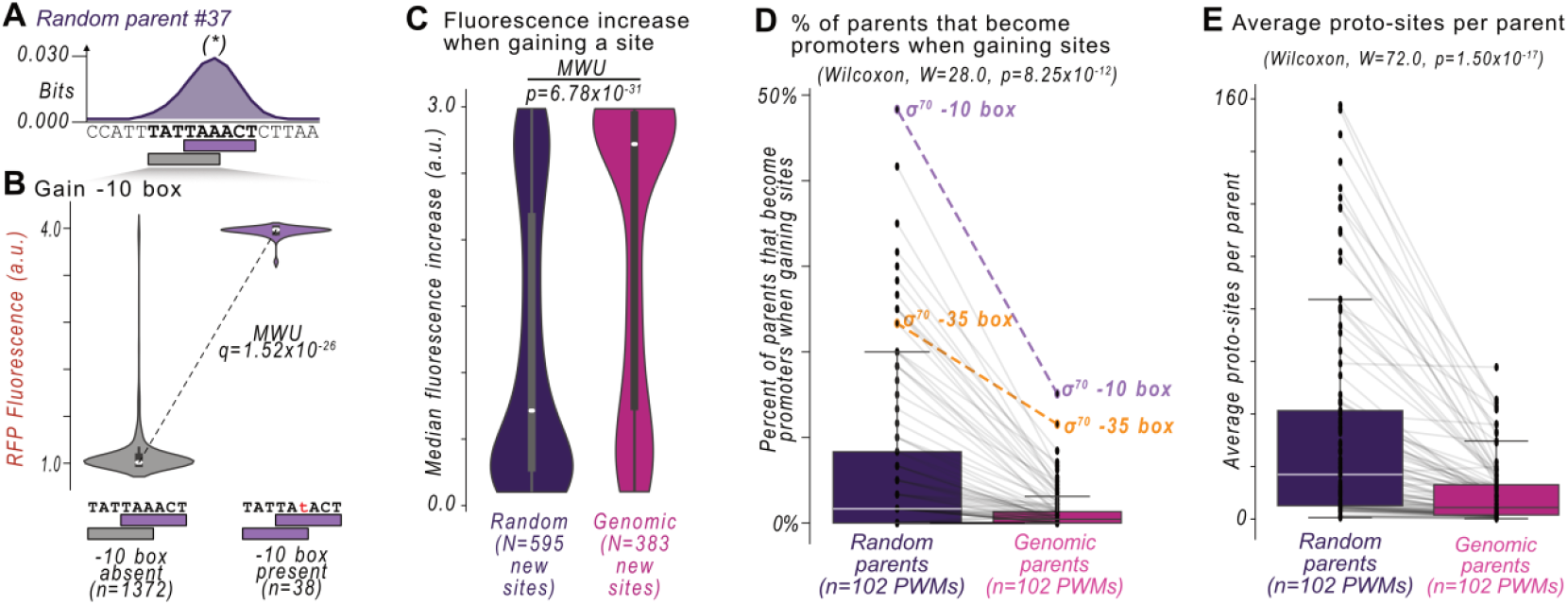
Promoters emerge less frequently in the genome because it harbors fewer proto-binding sites. (**A**) Top: Mutual information *I*_*i*_ *(b,j)* (vertical axis) at each position *i* (horizontal axis) between nucleotide identity *(j* = *A, T, C or G)* and fluorescence *(f* = *1*.*0 – 4*.*0 a*.*u*.) in information theoretical units (bits) for *random parent #37*. The dashed line indicates the 0.0025 bit threshold above which we call a mutual information peak a “hotspot” (asterisk). Bottom: outlined rectangles and bold sequences correspond to position-weight matrix (PWM)-predicted putative sites for the sigma 70 −10 box (purple) and a region of interest (gray). **(B)** The fluorescence scores of daughter sequences with /without a −10 box in the region of interest from (A). We test the null hypothesis that there is no difference in fluorescence between the two groups (two-tailed Mann-Whitney U [MWU] test), and display corrected q-values using the Benjamini-Hochberg procedure ^42^ (two-tailed MWU test, q=1.52×10^−26^). **(C)** The median fluorescence increase when gaining a putative binding site creates promoter activity. Purple = random parents, magenta = genomic parents (two-tailed MWU test, p=6.78×10^−31^). **(D)** The percentage of parent sequences that become promoters when gaining one of 102 PWM-predicted sites (N=60 random parents and 225 genomic parents). Each data point corresponds to a unique PWM. Purple = random parents, magenta = genomic parents. Grey lines pair each PWM between the random and genomic parents. We test the null hypothesis that there is no difference between the median percentages, using a Wilcoxon signed-rank test (W=28, p=8.25×10^−12^). **(E)** Analogous to (D) but for the average number of proto-sites per parent instead of the percentage of parent sequences (Wilcoxon signed-rank test, W=72, p=1.50×10^−17^). See **Source Data**.

This analysis revealed 595 and 383 activating sites in the random and genome parents, respectively, or ~9.9 new sites per random parent (595/60 parents) vs ~1.7 per genomic parent (383/225 parents). The increase in gene expression from gaining new sites was on average ~1.7 times higher (+1.97 a.u. vs +1.16 a.u.) in the genomic parents compared to the random parents (**Fig 3C**) (two-tailed MWU test, p=9.26×10^−31^). Simply put, genomic parents are less likely to become promoters by gaining binding sites. However, the de-novo promoters from genomic DNA are stronger than the de-novo promoters from random parents.

To understand how each of the 102 binding sites contributes to de-novo promoter emergence, we calculated the percentage of random and genomic parents that can become promoters when gaining each site. Overall, these percentages are lower in the genomic parents than in the random parents (**Fig 3D**) (Wilcoxon signed-rank test, p=8.25×10^−12^). For example, ~48% of random parents (29/60) acquire promoter activity by gaining a σ^70^ −10 box, while approximately three times fewer genomic parents do (~15% of 34/225). Similarly, ~23% of random parents (14/60) acquire promoter activity by gaining a σ^70^ −35 box, while almost two times fewer genomic parents do (~12% of 26/225). See **Source Data** for the frequencies of all 102 binding sites.

One possible explanation for this difference is that the genomic parents have fewer *proto-sites*, i.e. sequences that resemble TF and σ binding sites and can be converted into such a site by mutation. To test this hypothesis, we searched the parent sequences for DNA sequences yielding a positive PWM score, but where that score falls below the classification threshold as a putative binding site (**Methods**). Indeed, relative to random parents, genomic parents contain almost four times (median ~16.9 vs ~4.4) fewer proto-sites for the tested TFs and σ-factors (**Fig 3E**) (Wilcoxon signed-rank test, p=1.50×10^−17^). See **Source Data** for all 102 binding sites.

In sum, our analysis shows that emergence hotspots in which point mutations create a σ or TF binding site that *activate* transcription can help to explain why promoters emerge less frequently in genomic DNA: Genomic DNA contains fewer proto-sites than random DNA. We next turn to hotspots in which point mutations destroy sites that *repress* transcription.

### Genomic DNA is de-repressed compared to random DNA

A paucity of proto-sites in genomic DNA cannot explain why promoters that emerge in genomic DNA are stronger (see **Fig 3C**). This leaves the possibility that the stronger promoters that emerge in genomic DNA result from the genomic DNA being de-repressed compared to random DNA (see **Fig 1D**). To test this hypothesis, we screened the daughter sequences for instances where the loss of a predicted binding site for one of the 93 TFs, on either DNA strand, lead to increasing fluorescence (**Methods**). For example, destroying a predicted IclR site in random parent *#30* increases fluorescence by +1.41 a.u. (two-tailed MWU test, q=3.27×10^−8^) (See **Fig S8A,B** for this and additional examples). We call sites whose destruction causes increased transcription *repressing sites*.

We identified 131 repressing sites in random parents. They occur in more than half of the random parents (~58%, 35/60). Conversely, genomic parents harbor almost five times fewer repressing sites (27 sites vs 131 sites, in only 11 of the 225 genomic parents, **Fig 4A**). Remarkably, destroying any one of these 27 repressing sites increases fluorescence on average almost 13 times more than destroying a repressing site in a random parent (**Fig 4B**, random median: +0.04 a.u., genome median: +0.51 a.u., two-tailed MWU test, p=2.55×10^−10^). In other words, although genomic sequences are ~5× less likely to be subject to repression, their repression is ~13× stronger. For brevity, genomic DNA is de-repressed compared to random DNA.

**Figure 4.**
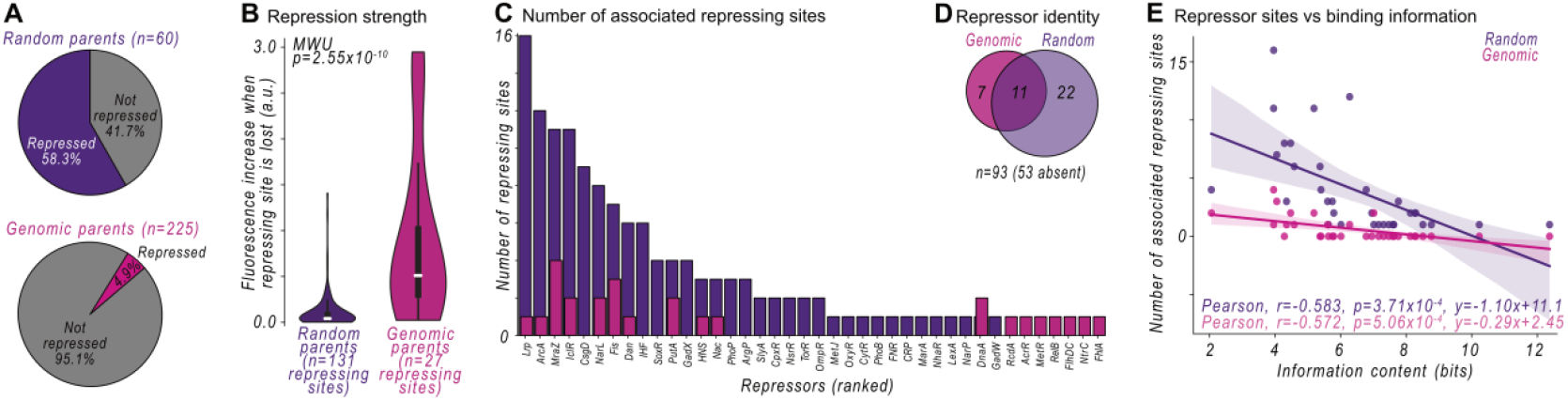
Genomic DNA is bound by fewer putative repressors, but repression is stronger. **(A)** A repressing site in a parent sequence is a site in which mutations that destroy a computationally predicted site result in increased fluorescence (**Methods**). We plot the percent of parents with a repressing site for the random (top, purple) and the genomic parents (bottom, magenta). **(B)** The median fluorescence increase in arbitrary units (a.u.) when a repressing site is destroyed in a random (purple) and genomic (magenta) parent (two-tailed Mann-Whitney U [MWU] test, p=2.55×10^−10^). **(C)** The number of putative repressor sites (ranked) for random (purple) and genomic (magenta) parents. **(D)** Venn-diagram indicating how many of the 93 tested transcription factors are predicted to bind to the random (purple) and genome (magenta) parents. **(E)** The number of repressing sites (vertical axis) vs their respective information content (horizontal axis, bits) for the random parents (purple) and the genomic parents (magenta). The solid line shows a best-fit linear regression line (shaded area: 95% confidence interval). See **Source Data**.

### Information theory can explain the greater de-repression of genomic DNA

Upon counting the number of repressing sites for specific transcriptional repressors in the random and genomic parents (**Fig 4C**), we found that 53 of the 93 tested repressors do not significantly repress transcription in the random or genome parents (**Fig 4D**, see **Fig S8** for examples). For the remaining 40 repressing TFs, 22 repress only the random parents, 7 repress only the genome parents, and 11 repress both the random and genome parents. In other words, the random parents are repressed by more unique TFs than the genomic parents (33 vs 18 repressors). We emphasize that repressing sites are not merely computationally predicted. Their loss is significantly associated with increasing transcription, and they overlap mutual information hotspots (see **Fig S8** for examples).

We hypothesized that the enrichment of repressor binding sites in random DNA is caused by chance alone, and by the binding specificity (or lack thereof) of the respective repressors. To this end, we compared repressor binding specificity (i.e., information content) vs the number of repressing sites in both genomic and random libraries. Not surprisingly, repressors with information-rich binding sites have fewer such sites, in both random and genomic DNA (**Fig 4E**, Pearson correlations. Random: r=-0.583, p=3.71×10^−4^; Genome: r=-0.572, p=5.06×10^−4^; see also **Fig 1D**). However, the slope *m* of the line of best fit is almost four times steeper for the random parents, and nearly flat in the genomic parents (m=-1.10 vs −0.29). This means that the dependency between the number of sites and information content is much weaker for genomic DNA than for random DNA. In other words, random chance can explain the abundance of repressor binding sites in the random library, but not in the genome. This abundance requires alternative explanations, such as natural selection.

## Discussion

To summarize our findings, we first compared the incidence of promoters in randomly synthesized DNA and genomic DNA (see **Fig 1**). Genomic DNA encodes ~1.3 times (~19.0% vs ~14.6%) more promoters than random DNA when we account for differences in AT-content, but also contains on average ~10 percent fewer (~16% vs ~26%) computationally predicted TF and σ factor binding sites than random DNA. Previous studies have made similar observations for σ^70^ sites^18,19^. We then compared how de-novo promoters emerge from random and genomic sequences upon mutation (see **Fig 2**). We found that promoters are ~3 times (median *Pnew* = 0.029 vs 0.092) less likely to emerge from genomic DNA than from random DNA. The reason is that genomic DNA encodes ~3.8 times fewer emergence hotspots (~0.4 hotspots vs ~1.5 hotspots), as a mutual information analysis of our experimental data reveals.

The paucity of emergence hotspots in genomic DNA has two explanations. First, the genome encodes fewer proto-sites (see **Fig 3**), sites that are easily converted into a promoter by mutation (~16.9 vs ~4.4 proto-sites per parent). This scarcity lowers the likelihood that mutations create new activating sites, leads to ~3.8 times fewer emergence hotspots, and ultimately makes promoters ~3 times less likely to emerge by mutations in genomic DNA. Second, the majority (~58%) of random parents are weakly repressed (see **Fig 4**), compared to a minority (~5%) of genomic parents that are strongly repressed. This difference may explain why promoters specifically inside genomic coding sequences are ~1.3 times more likely to be promoters compared to random DNA.

The scarcity of activating and repressing binding sites in the genome may be a product of natural selection. For example, selection may remove promoter-like sequences and TF binding sites to prevent the spontaneous emergence of promoters^27–29^. It may also affect binding sites indirectly by acting on other traits associated with gene expression^43^. For example, it may reduce conflicts between transcription and replication during cell division^44^, or it may be the reason why lowly expressed factors are sequestered at key binding sites^45–47^ elsewhere in the genome. To disentangle these different roles of natural selection remains an exciting task for future work.

Regardless of the selective pressures involved, the scarcity of activating proto-sites and repressing sites in genomic DNA will influence the evolution of new promoters in the genome. First, a paucity of repression biases the genome towards producing strong de-novo promoters (see **Fig 3C**). We speculate that this could lead to larger changes in fitness upon emergence, which would result in selection (rather than genetic drift) to act on their activity^26^. Second, because it harbors few activating proto-sites, the genome is biased against creating de-novo promoters. By extension, this bias would also constrain de-novo gene birth^16^.

In response to these biases, bacteria may depend on “domesticating”^48,49^ mobile and horizontally-transferred DNA as substrates for producing novel promoters^17,24^. This phenomenon has been observed during long term experimental evolution^17^, and in the evolution of antibiotic resistance in nature^50^. It is also similar to how enhancers emerge in eukaryotes^51–53^. For instance, most primate-specific TF binding sites have originated from transposable elements^54^.

Emergent promoters can create new transcripts for de-novo genes^16,17^, or transcribe previously inactive genes^13–15^. Either of these cases can potentially confer a selective advantage. We show here that the *E. coli* genome biases the *likelihood* of promoters emerging, *where* they emerge, *how* they emerge (by gaining or losing binding sites), *which* TFs they use, and the *strength* of their activity. By extension, such biases will influence the emergence of evolutionary adaptations and innovations.

## Methods

### Position-weight matrices (PWMs)

To generate position-weight matrices (PWMs), we acquired lists of putative binding sites for transcription factors (TFs) from RegulonDB^38^ and individual publications for σ factors^55–58^. Using the `motifs` package from `Biopython` (v1.79)^59^, we created position weight matrices for these factors, assuming a uniform background frequency of 25% for each nucleotide. We then classified any one query sequence as a binding site if its respective PWM score (in bits) was greater than or equal to the natural logarithm of the PWM’s information content (in nats). This threshold is also referred to as the Patser Threshold^60^. In **Fig 3E**, we also classified sequences as proto-binding sites if the PWM score was greater than 0.00 bits, but less than this Patser Threshold.

### Creating the wild-type genomic library

See **Fig S9** for an overview of how we created the wild-type genomic library. To isolate genomic DNA (gDNA), we inoculated a culture of DH5α electrocompetent *E. coli* cells (Takara, Japan, product #9027) and allowed the cells to grow overnight with shaking at 230 rpm at 37°C. The following day, we extracted the gDNA using a Wizard kit (Promega USA, product #A1120) according to the manufacturer’s instructions, eluting the final product in 50 uL of water (**Fig S9A**). We estimated the gDNA concentration using a Nanodrop One spectrophotometer (Thermo Scientific, USA) as ~10 ng in a 50 uL total volume.

We sheared the gDNA into fragments using a focused ultrasonicator (Covaris E220), with the following parameters: 1) Duty factor: 10%; 2) Peak Incident Power (W) 175; 3) Cycles per Burst 200; 4) Time (seconds) 430, as described on the manufacturer’s website (https://www.covaris.com/e220-focused-ultrasonicator-500239)

We separated the fragmented gDNA using agarose gel electrophoresis with a 3% agarose gel, and manually excised DNA between 100 and 200 bp in length. We purified this DNA using a Qiagen QIAquick Gel Purification Kit (Qiagen, Netherlands, product #28706) according to the manufacturer’s instructions. To repair any DNA damage caused by sonication, we used a NEBNext repair kit (NEB, USA, product #M6630) to create blunt ends. We did this by adding 270 ng of the sheared DNA (18 uL), 2.5 uL of the provided buffer, 1.25 uL of the provided enzymes, and 3.25 uL of water (25 uL total volume). We incubated this reaction for 30 minutes at 20°C in a C1000 Touch Thermal Cycler (Bio-Rad, USA) with the thermal cycler lid temperature at 30°C. We purified the reaction product using a QIAquick PCR Purification Kit (Qiagen, Netherlands, product #28104) according to the manufacturer’s instructions.

We next ligated an Adenine (`A`) base to the 3′-ends of each fragment using the NEBNext dA-Tailing Module (NEB, USA, product #E6053) (**Fig S9B**) by combining 20 uL of blunt-end DNA (21 ng/uL), 2.5 uL of buffer, 1.5 uL of the Klenow fragment, and 1.0 uL of H_2_O (25 uL total volume), and incubating the reaction at 37°C for 35 minutes in a C1000 Touch Thermal Cycler with the lid heated to 45°C. We purified this product with a QIAquick PCR Purification Kit (Qiagen, Netherlands, product #28104) according to the manufacturer’s instructions.

To create upstream and downstream double-stranded DNA sequences homologous to our expression vector, we synthesized and annealed two sets of complementary oligonucleotides (IDT, Coralville USA). For each double-stranded DNA, one of the oligonucleotides included a 5′ phosphorylation modification for downstream ligation (See **Supplementary Data 7** for primer sequences for oligo1-4). We annealed complementary oligonucleotides by incubating them in an equimolar ratio at 95°C for two minutes in a thermocycler (C1000 Touch Thermal Cycler, Bio-Rad, USA), and subsequently decreasing the temperature by increments of 5°C every 90 seconds, until a final temperature of 25°C had been reached.

To ligate the double-stranded DNA upstream and downstream of the adenylated genomic DNA, we used a T4 ligase master mix (NEB, USA, product #M0202), with 3 uL of 10× T4 buffer, 1 uL of the upstream annealed adaptor (100 uM), 1 uL of the downstream annealed adaptor (100 uM), 20 uL of the adenylated genomic DNA (16 ng/uL, 320 ng total), 1.5 uL of T4 Ligase, and 3.5 uL of H_2_O. We incubated this reaction mix at room temperature for two hours, and heat-inactivated the reaction at 65°C for 10 minutes. Following the ligation, we excised the product using agarose (1%) gel electrophoresis, and purified the isolated DNA using a QIAquick Gel Purification Kit (Qiagen, Netherlands, product #28706) according to the manufacturer’s instructions.

We performed a polymerase chain reaction (PCR) to amplify the ligated products using Q5 high-fidelity polymerase (NEB, USA, product #M0491) according to the manufacturer’s instructions, and with primers “pMR1_insert_forward” and “pMR1_insert_reverse” provided in **Supplementary Data 7**. We incubated the reaction for 30 cycles (annealing during each cycle at 55°C for 30 seconds and extending primers at 72°C for 30 seconds) using a thermal cycler (C1000 Touch Thermal Cycler, Bio-Rad, USA). We then separated the PCR products on a 1% agarose gel, excised the product between 200-500 bp, and purified the gel product using a QIAquick Gel Purification Kit (Qiagen, Netherlands, product #28706). We cloned the genomic library into the plasmid MR1 (pMR1) using Gibson Assembly (**Fig S9C**), and transformed the products into DH5α cells as described in the subsection ***Molecular Cloning and Transformations***.

### Creating the wild-type random library

To create the wild-type random library, we synthesized a 150-Nmer sequence with flanking sequences homologous to our reporter vector (pMR1) (IDT, Coralville USA). See **Supplementary Data 7** for the homologous DNA sequences. Using Q5 high-fidelity polymerase (NEB, USA, product #M0491), we amplified the fragment according to the manufacturer’s instructions, and with the primers shown in **Supplementary Data 7**. We incubated the PCR reaction for 30 cycles (annealing in each cycle at 55°C for 30 seconds, and extending the primers at 72°C for 30 seconds) using a thermal cycler (C1000 Touch Thermal Cycler, Bio-Rad, USA). We then separated the PCR products on a 1% agarose gel, excised DNA of ~200-300 bps with a scalpel, and purified the gel product using a QIAquick Gel Purification Kit (Qiagen, Netherlands, product #28706) according to the manufacturer’s instructions. We cloned the library into the pMR1 plasmid using Gibson Assembly, and transformed the resulting plasmid library into DH5α cells as described in the ***Molecular Cloning and transformations*** section.

### Molecular cloning and transformations

To clone inserts into the pMR1 dual reporter plasmid ^35^, we used Q5 polymerase (NEB, USA, product #M0491) to make linearized copies of the pMR1 plasmid flanking the BamHI (`GGATCC`) and EcoRI (`GAATTC`) cut sites. To this end, we used primers “pMR1_vector_forward” and “pMR1_vector_reverse” listed in **Supplementary Data 7** and the following reagents: 20 uL Q5 buffer, 2 uL dNTPs (Thermo Scientific, USA, product #R0191), 10 uM forward primer, 10 uM reverse primer, 1 uL of the insert, 1 uL Q5 polymerase, 71 uL H_2_O. We performed the cloning reaction in a thermal cycler (C1000 Touch Thermal Cycler, Bio-Rad, USA), for 30 cycles of annealing at 55°C for 30 seconds, and extending at 72°C for 2 minutes and 30 seconds. We separated the cloning products on a 1% agarose gel, and extracted the DNA band of interest using a scalpel. To purify the gel product, we used a QIAquick Gel Purification Kit (Qiagen, Netherlands, product #28706) according to the manufacturer’s instructions, and eluting the DNA in 50 uL of H_2_O.

We then used NEBuilder (Gibson Assembly) (New England Biolabs, USA, product #E2621) to clone inserts into the linearized plasmid pMR1. To this end, we added 100 ng of pMR1, 5 uL of the NEBuilder mastermix, 24 ng of insert, and H_2_O to a total volume of 10 uL. We incubated this cloning reaction for 1 hour at 50°C. Subsequently, we transformed 2 uL of the cloned product into 100 uL of DH5α electrocompetent cells (Takara, Japan, product #9027) using an electroporator (Bio-Rad, USA, MicroPulser). Following electroporation, we allowed the cells to recover in 1 mL of SOC (Super Optimal Broth with Catabolite Repression Medium, Merck, product #CMR0002) for 1.5 hours, incubating at 37°C and shaking at 230 rpm. We plated 5 uL of the transformed culture onto a Luria Broth (LB) agar plate supplemented with 100 ug/mL of chloramphenicol (cm). We added 2 mL of LB-cm to the remaining 995 uL of transformed product, and cultured the cells overnight.

The following morning, we counted the number of colonies on the agar plate to estimate the total number of unique inserts in the library. We then stored 1 mL of the liquid library culture with 667 uL 60% glycerol at −80°C for later use. With the remaining 2 mL of the liquid culture, we isolated the library using a QIAprep Spin Miniprep Kit (Qiagen, Germany, product #27104).

### Defining fluorescence bins for Sort-Seq

We defined the boundaries for G1-G4, R1-R4, GFP-on/off, and RFP-on/off bins using the fluorescence readouts of three control reporter plasmids (**Fig S10**). The first was a negative control consisting of the pMR1 plasmid backbone without an insert. The second was a positive control for green fluorescence, which contains the *bba_J23110* promoter inserted into pMR1 oriented towards the green fluorescent protein (GFP) coding sequence. The third was a positive control for red fluorescence, which contains the *bba_J23110* promoter inserted into pMR1 oriented towards the red fluorescent protein (RFP) coding sequence.

For the green fluorescence bins, we first defined the lower boundary of G1 (no green fluorescence) as the lowest measured green fluorescence in the negative control (pMR1 with no insert) and the lowest measured green fluorescence for the RFP positive control. We use a lower bound to avoid counting and isolating debris such as salts and cell waste. To define the upper boundary of G1, we used the highest measured green fluorescence in the negative control, and the highest measured green fluorescence in the RFP positive control. To define the boundaries of R1 (no red fluorescence), we repeated this procedure, but switched the positive RFP control with the GFP positive control.

To define the lower boundaries of G4 and R4 (strong green and red fluorescence, respectively), we used the mean fluorescence of the respective (green or red) positive control. We did not define an upper bound for bins G4 and R4. We note that the GFP distribution for the GFP positive control was bimodal, where the lower (left) peak likely corresponds to cells not harboring the insert (see **Fig S10B**). For this reason, we take the mean fluorescence of the second (right) peak to define the lower bound of G4 (**Fig S10B**).

To define the boundaries of G2 and G3, we divided the lower boundary of G4 and the upper boundary of G1 in half, such that the lower bound of G3 becomes the upper bound of G2. Similar for R2 and R3, we divided the lower boundary of R4 and the upper boundary of R1 in half, such that the lower bound of R3 becomes the upper bound of R2. See **Fig S10** for an overview of the controls and bin boundaries. See also subsections ***Sort-Seq (wild-type Random and Genomic Libraries)*** and ***(Sort-Seq mutagenesis libraries)*** for sorting details.

For the genome mutagenesis library, we sorted cells into the four bins GFP-ON, GFP-OFF, RFP-ON, and RFP-OFF, as opposed to the eight bins just described (G1-G4 and R1-R4). GFP-OFF is analogous to G1 and RFP-OFF is analogous to R1. The lower bound of GFP-ON is analogous to the lower bound of G2, and does not have an upper bound. Likewise, for RFP-ON the lower bound is analogous to the lower bound of R3, and does not have an upper bound. See **Fig S10 and *Sort-Seq (mutagenesis libraries)*** subsection for sorting details.

### Sort-Seq (wild-type random and genomic libraries)

We inoculated 100 uL of the random library or genomic library into separate cultures of 10 mL Luria Broth (LB) supplemented with 100 ug/mL of chloramphenicol, and incubated the two cultures at 37°C overnight with shaking at 230 rpm. The following morning (ca. 16 hours later), we washed and resuspended the bacteria with Phosphate Buffer Saline (PBS). We then sorted the random and genomic libraries independently into 4 green fluorescence bins (G1, G2, G3, and G4) and 4 red fluorescence bins (R1, R2, R3, and R4), using a cell sorter (BD-Biosciences, FACSAria III) (see subsection ***Defining fluorescence bins for Sort-seq*** for binning procedure) as follows: For the wild-type random library (day 1), we sorted 1’440’000 cells into G1, 180’000 cells into G2, 120’000 cells into G3, 60’000 cells into G4, 4’320’000 cells into R1, 540’000 cells into R2, 360’000 cells into R3, and 180’000 cells R4. For the wild-type genomic library (day 1), we sorted 1’200’000 cells into G1, 150’000 cells into G2, 100’000 cells into G3, 864 cells into G4, 1’200’000 cells into R1, 150’000 cells into R2, 33’647 cells into R3, and 17’101 cells into R4.

After sorting (8 bins for the 2 libraries, 16 bins in total) we added 1 mL of SOC (Super Optimal Broth with Catabolite Repression Medium, Merck, product #CMR0002) to each culture, and incubated the cultures at 37°C (with shaking at 230 rpm) for two hours to help the bacteria recover. We then supplemented each culture with an additional 2 mL of LB and 0.3 ug of chloramphenicol (3mL total volume). We incubated the 8 cultures for both libraries overnight at 37°C with shaking at 230 rpm. The following morning (ca. 16 hours later), we repeated the sorting and recovery procedure, this time, re-sorting each culture into its corresponding bin as the day before, in triplicates, as follows: For the wild-type random library (day 2), we re-sorted 1’080’000 total cells into G1 (divided into 3 equal-volume replicates [r1, r2, r3], ~360’000 cells per replicate), 104’222 total cells into G2, 25’678 total cells into G3, 45’000 total cells into G4, 1’080’000 total cells into R1, 135’000 total cells into R2, 90’000 total cells into R3, and 45’000 total cells into R4 (3 replicates, 8 bins, 24 final cultures). For the wild-type genomic library (day 2), we re-sorted 3’600’000 total cells into G1, 450’000 total cells into G2, 300’000 total cells into G3, 30’000 total cells into G4, 3’600’000 total cells into R1, 450’000 total cells into R2, 100’941 total cells into R3, and 51’303 total cells into R4 (3 replicates, 8 bins, 24 final cultures).

We allowed the bacteria in each of the 24 cultures to recover for two hours in SOC before adding an additional 2 mL of LB with 0.3 ug of chloramphenicol. We then incubated the cultures overnight at 37°C at 230 rpm. The following morning (ca. 16 hours later), we stored 1 mL of each library culture with 667 uL 60% glycerol at −80°C. With the remaining 2 mL of the culture, we isolated the plasmids from each culture using a QIAprep Spin Miniprep Kit (Qiagen, Germany, product #27104).

### Creating the mutagenesis libraries

We inoculated 100 uL of the random library and the genome library into separate cultures of 10 mL Luria Broth (LB) supplemented with 100 ug/mL of chloramphenicol, and incubated the two cultures at 37°C overnight at 230 rpm. The following morning (ca. 16 hours later), we washed and resuspended the bacteria with Phosphate Buffer Saline (PBS). Using a cell-sorter (BD Biosciences, FACSAria III), we isolated 2,000,000 and 1,000,000 bacteria that collectively fluorescence in both bin G1 (no GFP fluorescence) and bin R1 (no RFP fluorescence) from the random library and the genomic library, respectively (two sorted cultures after sorting). We allowed the sorted bacteria to recover in 1 mL of SOC (Super Optimal Broth with Catabolite Repression Medium, Merck, product #CMR0002) for two hours before we added an additional 2 mL of LB and 0.3 ug of chloramphenicol (to 3mL of total volume). We incubated the two cultures overnight at 37°C with shaking at 230 rpm. The following morning (ca. 16 hours later), we repeated the sorting and recovery procedure for these cultures, this time isolating 459’185 bacteria that sort into G1 and R1 from the negative random library culture, and 1’611’091 bacteria that sort into G1 and R1 from the negative genomic library culture. We refer to these cultures as the “negative” random (genomic) cultures.

We plated the negative cultures onto LB-Agar petri dishes supplemented with chloramphenicol (cm, 100 ug/mL). We then picked individual colonies using pipette tips, resuspending each colony in 10 uL of H_2_O. We selected 96 colonies from the negative random culture, and 396 colonies from the negative genomic cultures. We then carried out individual error-prone polymerase chain reactions (EP-PCRs) for the DNA in each colony. For each EP-PCR, we added 6.9 uL of H2O, 2 uL of 5x GoTaq reaction buffer, 0.2 uL of GoTaq polymerase (Promega, USA, product #M3001), 0.1 uL of Primer “pMR1_insert_forward”, 0.1 uL of Primer “pMR1_insert_reverse”, 0.2 uL of dNTPs (Thermo Fischer, USA, product #R0191), and 0.2 uL of MnCl_2_ at 15 mMol (for mutagenesis) to a reaction “master-mix”. For each EP-PCR, we added 9 uL of this master-mix to 1 uL of the resuspended colony. We then performed the reactions in a thermal cycler (C1000 Touch Thermal Cycler, Bio-Rad, USA) for 30 cycles of annealing at 55°C for 30 seconds and extending at 72°C for 30 seconds. We pooled the products of the random EP-PCR into one test tube and the genomic EP-PCR into another test tube. We separated the pooled reaction products electrophoretically on a 1% agarose gel, and extracted the DNA bands of interest using a scalpel. To purify the gel products, we used a QIAquick Gel Purification Kit (Qiagen, Netherlands, product #28706) according to the manufacturer’s instructions. We separately cloned and transformed the two mutagenized insertion libraries into plasmid MR1 (pMR1) and DH5α electrocompetent cells as described in subsection ***Molecular cloning and transformations***.

### Sort-Seq (Mutagenesis libraries)

We inoculated 100 uL of the random mutagenesis library and the genomic mutagenesis library into two separate cultures of 10 mL Luria Broth (LB) supplemented with 100 ug/mL of chloramphenicol, and incubated the two cultures at 37°C overnight at 230 rpm. The following morning (ca. 16 hours later), we washed and resuspended the bacteria in Phosphate Buffer Saline (PBS).

We sorted the genomic mutagenesis library into 2 green fluorescence bins (GFP-ON and GFP-OFF) and 2 red fluorescence bins (RFP-ON and RFP-OFF). In addition, we sorted the random mutagenesis library into 4 green fluorescence bins (G1, G2, G3, and G4) and 4 red fluorescence bins (R1, R2, R3, and R4) (see subsection ***Defining fluorescence bins for Sort-seq*** for binning procedure). GFP-ON and GFP-ON are respectively analogous to G1 and R1. However, we combined the read counts (post-sequencing) from bins R2-R4 and G2-G4 to make these combined bins analogous to RFP-ON and GFP-ON, respectively, as described in the subsection ***Processing mutagenesis library sequencing results***.

For the random mutagenesis Library (day 1), we sorted 1’000’000 cells into G1, 70’826 cells into G2, 11’129 cells into G3, 12’576 cells into G4, 1’000’000 cells into R1, 324’742 cells into R2, 52’550 cells into R3, and 24’276 cells into R4. For the genomic mutagenesis library (day 1), we sorted 2’000’000 cells into GFP-OFF, 5’754 cells into GFP-ON, 2’000’000 cells into RFP-OFF, and 193’330 cells into RFP-ON.

We allowed the sorted bacteria in each bin to recover in 1 mL of SOC (Super Optimal Broth with Catabolite Repression Medium, Merck, product #CMR0002) for two hours before supplementing each culture with an additional 2 mL of LB and 0.3 ug of chloramphenicol (3mL total volume). We incubated the cultures overnight at 37°C, with shaking at 230 rpm. The following morning (ca. 16 hours later), we repeated the sorting and recovery procedure, this time re-sorting each culture into its corresponding bin as follows: For the random mutagenesis library (day 2), we re-sorted 3’000’000 total cells into G1, 200’013 total cells into G2, 250’000 total cells into G3, 375’000 total cells into G4, 3’000’000 total cells into R1, 1’500’000 total cells into R2, 750’000 total cells into R3, and 375’000 total cells into R4 (8 bins, 8 cultures, no replicates). For the genomic mutagenesis library (day 2), we re-sorted 6’000’000 total cells into GFP-OFF, 360’000 total cells into GFP-ON, 6’000’000 total cells into RFP-OFF, and 1’960’687 total cells into RFP-ON (4 bins, 4 cultures, no replicates).

We allowed the sorted cultures to recover for two hours in SOC before adding an additional 2 mL of LB with 0.3 ug of chloramphenicol, and incubating the cultures overnight at 37°C, with shaking at 230 rpm. The following morning (ca. 16 hours later), we stored 1 mL of each culture with 667 uL 60% glycerol at −80°C. With the remaining 2 mL of the culture, we isolated the plasmids from each culture using a QIAprep Spin Miniprep Kit (Qiagen, Germany, product #27104) for *Illumina* sequencing.

### Illumina sequencing

From the plasmids isolated from each sorted library bin, we used a polymerase chain reaction (PCR) to amplify the plasmid inserts for next-generation sequencing. Specifically, for each PCR, we mixed 20 uL Q5 buffer (NEB, USA, product #4091), 2 uL dNTPs (Thermo Scientific, USA, product #R0191), 10 uM reverse primer “Illumina_reverse_primer”, 1 uL of the insert, 1 uL Q5 polymerase, and 71 uL H_2_O. In addition, we added to each reaction mix a multiplexing forward primer (10 uM) with a unique barcode for each fluorescence bin and replicate. See **Supplementary Data 7** for a list of primers and barcodes. We performed the reaction in a thermocycler (C1000 Touch Thermal Cycler, Bio-Rad, USA), with 30 cycles of annealing at 55°C for 30 seconds and extending at 72°C for 2 minutes and 30 seconds. We separated the PCR products on a 1% agarose gel, and extracted the DNA band of interest using a scalpel. To purify the gel product, we used a QIAquick Gel Purification Kit (Qiagen, Netherlands, product #28706) according to the manufacturer’s instructions. We pooled the PCR products and sequenced them using Illumina paired-end read sequencing (Eurofins GmbH, Germany) on an Illumina NovaSeq 6000 (Illumina, USA) with an S4 flow cell and paired-end 150 bp runs.

### Processing wild-type library sequencing results

For the wild-type libraries, we merged the sequencing reads into paired-end reads using Flash2 ^61^. We oriented each paired-end read based on the flanks of the read, and categorized the read to its respective bin and replicate number using the unique forward primer index. To map the genomic fragments to their respective genomic coordinates, we used bowtie2 (v2.3.5.1) to create an index map of the E. coli genome, and mapped the fragments to these indices using samtools (v1.6).

We excluded reads from further analysis, if they 1) were not present at least once in the unsorted library, 2) were not present in at least four separate bins, 3) were shorter than 100 bp or longer than 200 bp, 4) did not map to the genome (for reads from the genomic library). **Supplementary Data 1 and 2** are csv files of the remaining (included) reads from the random wild-type library and the genomic wild-type library, respectively, together with their fluorescence scores.

### Processing mutagenesis library sequencing results

For the mutagenesis libraries, we also merged the sequencing reads into paired-end reads using Flash2 ^61^. We oriented each paired-end read based on the flanks of the read, and categorized the read by its respective bin and replicate number using the unique forward primer index.

As described in subsection ***Sort-Seq (Mutagenesis Libraries)***, we had sorted the random mutagenesis library into eight fluorescence bins (G1-G4 and R1-R4), while we had sorted the genomic mutagenesis library into only four fluorescence bins (GFP-OFF, GFP-ON, RFP-OFF, and RFP-ON). G1 and R1 are analogous to GFP-OFF and RFP-OFF, while the sum of G2, G3, and G4 is analogous to GFP-ON, and the sum of R2, R3, and R4 is analogous to RFP-ON (see ***Defining fluorescence bins for Sort-seq*** subsection for binning procedure and **Fig S10**). To compare the results from the two mutagenesis libraries, we thus pooled the reads from G2, G3, and G4 to constitute the corresponding GFP-ON bin. Similarly, we pooled the reads of R2, R3, and R4 to constitute the corresponding RFP-ON bin.

To map the daughter (mutagenized) sequences to their respective parents (wild-type), we partitioned the daughters into groups based on their length in base pairs. For each group of sequences with the same length, we represented the Hamming distance between every sequence pair as a contingency matrix. From the contingency matrix, we identified clusters of sequences with a Hamming distance of no more than 10 mutations. From each cluster, we then identified the parent sequence as the consensus DNA sequence of the cluster.

We excluded a parent and its daughters from the dataset if 1) we could not find the parent sequence in the respective wild-type library, 2) there were less than 500 daughters per parent, 3) the parent and daughter sequences were shorter than 100 bps or longer than 200 bp. The resulting filtered sequence data sets are contained in **Supplementary Data 3** for the random mutagenesis library and **Supplementary Data 4** for the genomic mutagenesis library.

### Calculating fluorescence scores

We calculated fluorescence scores (*fluor*_*r*_) for each technical replicate (r1, r2, r3) and fluorophore (GFP and RFP) using **equation (1):**

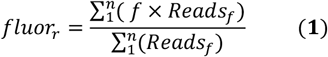

When this equation is applied to the wild-type libraries, the index f represents integer-encoded fluorescence bins (f=1, 2, 3, 4 for G1, G2, G3, G4 or R1, R2, R3, R4), and *Reads*_*f*_ is the number of reads within each fluorescence bin f. For each sequence, we computed the average fluorescence score across replicates using **equation (2)**:

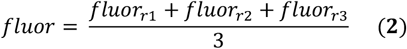

We do not have replicates for the mutagenesis libraries and therefore do not take the mean of the replicates for the fluorescence scores. Therefore *fluor* = *fluor*_*r*_ for the mutagenesis libraries.

### Mutual information

For each parent sequence, we quantified the mutual information *I*_*i*_ between the nucleotide identity *b* at position *i* (where 1 ≤ *i* ≤ the length of the parent sequence) in its corresponding daughter sequences and their associated fluorescence values. We performed this computation using **equation (3)**, following the method outlined in ref ^8^.

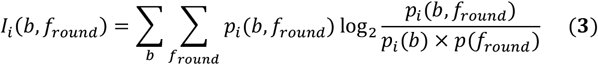

Here, b corresponds to all possible nucleotides (*b* = *A, T, C, G)*, and *f*_*round*_ corresponds to the daughter fluorescence scores *fluor* rounded to the nearest integer, yielding four discrete fluorescence levels (*f*_*round*_=1, 2, 3, and 4 a.u.) (see ***Processing sequencing results*** for calculating fluorescence scores). In other words, *f*_*round*_ is a discretized version of the continuous fluorescence score *fluor* defined in ***Processing sequencing results***.

The quantity *p*_*i*_*(b)* denotes the relative frequency of A, T, C, or G at position (*i)* in the daughter sequences; *p(f*_*round*_) is the relative frequency of rounded fluorescence scores being equal to 1, 2, 3, and 4 a.u.; and *p*_*i*_*(b, f*_*round*_) is the corresponding joint frequency.

We visualize mutual information hotspots – regions of high mutual information – by smoothening mutual information values as a function of position *i* with the `scipy` package `ndimage` Gaussian Filter (parameter alpha=2). See mutual information values in **Source Data**. From these smoothened data, we identified mutual information peaks at each position *i*, such that *I*_*i*_ *(b, f*_*round*_) > 0.0025 bits, and *I*_*i*−*1*_*(b, f*_*round*_ ) < *I*_*i*_ *(b, f*_*round*_) > *I*_*i*+*1*_*(b, f*_*round*_).

To confirm that our finding of random parents containing more hotspots than genomic parents (see **Fig 2D**) is not a byproduct of our thresholding and smoothing procedures, we repeated this analysis using three different thresholds (0.000, 0.0025, and 0.005 bits) and four different smoothing alpha parameters (0, 1, 2, and 3) (see **Fig S6**).

### Identifying significant associations between mutations creating or destroying sites

To identify regions within each parent sequence where binding-site creating (destroying) mutations are associated with increasing fluorescence, we performed the following computational screen for each parent sequence of length L bps. We moved a sliding window of length X—corresponding to the length (X) of the tested position weight matrix (PWM) along each parent sequence in one bp increments (from position 1 to position L-X). At each position, we scanned all corresponding mutant daughter sequences for the presence or absence of a PWM-predicted binding site, as defined in the subsection ***Position Weight Matrices (PWMs)***. We then grouped the corresponding fluorescence scores of daughter sequences with the binding site as “positives,” and those lacking the site as “negatives.”

If both the “positive” and “negative” group contained more than 10 fluorescence scores, we tested whether binding site presence was associated with higher fluorescence using a two-sided Mann–Whitney U test (via `mannwhitneyu` in `scipy.stats`). We applied this procedure at each window position, on both DNA strands, for all parent sequences, and for both green and red fluorescence measurements. To account for multiple hypothesis testing, we applied the Benjamini–Hochberg procedure ^42^ with a false discovery rate of 0.05, and considered regions with p-value corrected q-values below 0.05 (q < 0.05) to be significant. **Supplementary Data 5** and **Supplementary Data 6** contain the results of this analysis for the random mutagenesis library and the genome mutagenesis library, respectively.

To specifically explore how mutations create binding sites (see **Fig 3**), we focused on sites that 1) were gained (not lost); 2) emerged on the same strand associated with the fluorescence increase; 3) had an associated fluorescence increase greater than or equal to 0.1 arbitrary units (a.u.).

To specifically explore how mutations destroy binding sites (see **Fig 4**), we focused on sites that 1) were lost; 2) occurred on either DNA strand associated with a fluorescence increase; 3) showed an associated fluorescence increase greater than 0.0 a.u. We performed these analyses for the 93 binding sites (PWMS) listed in **Supplementary Data 5 and 6**. We did not analyze associations where losing a predicted σ factor binding site increases fluorescence, because our previous work demonstrates that such sites are false positives confounded by the creation or destruction of other predicted sites^23^.

## Data availability

The raw sequencing files from this study are available in the Sequence Read Archive (SRA) with the following accession code: PRJNA1304939: http://www.ncbi.nlm.nih.gov/bioproject/1304939 and BioSample accession numbers: SAMN50577760, SAMN50577761, SAMN50577762, and SAMN50577763.

We also provide the processed data in the GitHub repository: https://github.com/tfuqua95/random-genomic. The **Source Data** are provided as a Source Data file.

File name: **Table S1**

Description: a data frame (Microsoft Excel spreadsheet) containing putative promoter sequences from RegulonDB^38^ and their respective matches to the genomic sequences in our dataset.

File name: **Supplementary Data 1**

Description: a data frame (csv format) containing the wild-type random DNA sequences and their respective fluorescence scores.

File name: **Supplementary Data 2**

Description: a data frame (csv format) containing the wild-type genomic DNA sequences and their respective fluorescence scores.

File name: **Supplementary Data 3**

Description: a data frame (csv format) containing the random parent mutagenesis library sequences and their respective fluorescence scores.

File name: **Supplementary Data 4**

Description: a data frame (csv format) containing the genome parent mutagenesis library sequences and their respective fluorescence scores.

File name: **Supplementary Data 5**

Description: a data frame (csv format) containing the associations between gaining or losing transcription factor or sigma factor binding sites and the change this has on fluorescence scores in the random parent sequences.

File name: **Supplementary Data 6**

Description: a data frame (csv format) containing the associations between gaining or losing transcription factor or sigma factor binding sites and the change this has on fluorescence scores in the genome parent sequences.

File name: **Supplementary Data 7**

Description: a data frame (Microsoft Excel spreadsheet) containing primer DNA sequences for polymerase chain reactions (PCRs) and molecular cloning.

**Supplementary Data 1-7, Table S1**, and **Source Data** are also available on the GitHub repository: https://github.com/tfuqua95/random-genomic

## Code availability

A Jupyter Notebook containing the Python scripts to carry out all analyses, an Anaconda environment with the relevant packages and their respective versions, as well as the **Source Data, Table S1**, and **Supplemental Data 1-7** are located at the GitHub repository: https://github.com/tfuqua95/random-genomic

## Acknowledgements

This research was supported by the Swiss National Science Foundation (grants 31003A_172887 and 310030_208174) and the European Research Council (Grant Agreement No. 739874). TF is supported by a fellowship from the European Molecular Biology Organization (ALTF 963-2021) and a University of Zurich Postdoc Grant (FK-23-120). We thank all members of the Wagner group for discussions, especially Baxter.

---

**Fig S1.**
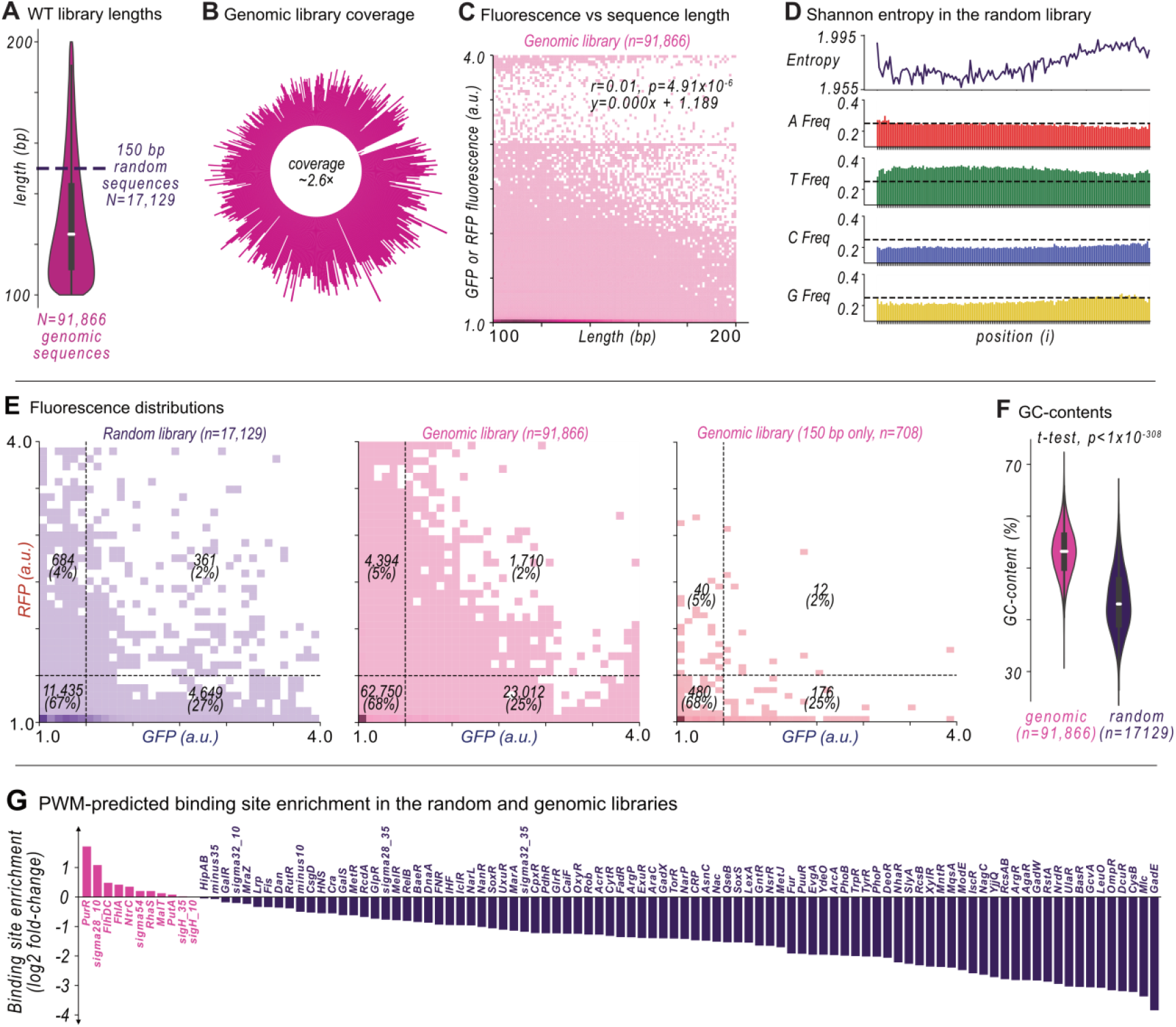
Wild-type library compositions. **(A)** The distribution of sequence lengths in the genomic library. The horizontal dashed line corresponds to the fixed 150 bp length of sequences in the random library. The white line shows the median, the gray box the interquartile range (IQR), whiskers span ±1 standard deviation, and the violin area reflects a kernel density estimate of the distribution. **(B)** A histogram of the coordinates of sequences in the genomic library. **(C)** Scatter plot of fluorescence scores (y-axis) vs sequence length (x-axis) of sequences in the genomic library. We tested the null hypothesis that there is no association between fluorescence and length. Although this association is significant, it is very weak (Pearson’s r = 0.01, p=4.91×10^−6^, n=91,866). The fitted linear equation on top of the panel was calculated using the method of least squares (line of best fit not shown). **(D)** Top: the Shannon Entropy of nucleotide identity at each position i in the random library. Bottom four panels: the frequency of each nucleotide (A, T, C, or G) at each position. The dashed horizontal line represents a frequency of 0.25. **(E)** Scatter plot of fluorescence scores (x-axis GFP, y-axis RFP) for sequences in the libraries (left: random library, middle: genomic library, right: genomic library with exactly 150 bp in length. Dashed lines at 1.5 a.u. indicate our threshold for promoter activity. **(F)** GC-content of the genome (magenta, left) vs the random (purple, right) libraries (two-tailed t-test, p<1×10^−308^). **(G)** For 102 position-weight matrices (PWMs) for transcription factors and sigma (σ) factors we plot the percentage of sequences in each library (purple: random, magenta: genome) that encode at least one putative binding site for each factor (See **Fig 1D**). We show the log_2_ fold-enrichment of these frequencies as a bar plot. See **Source Data**.

**Fig S2.**
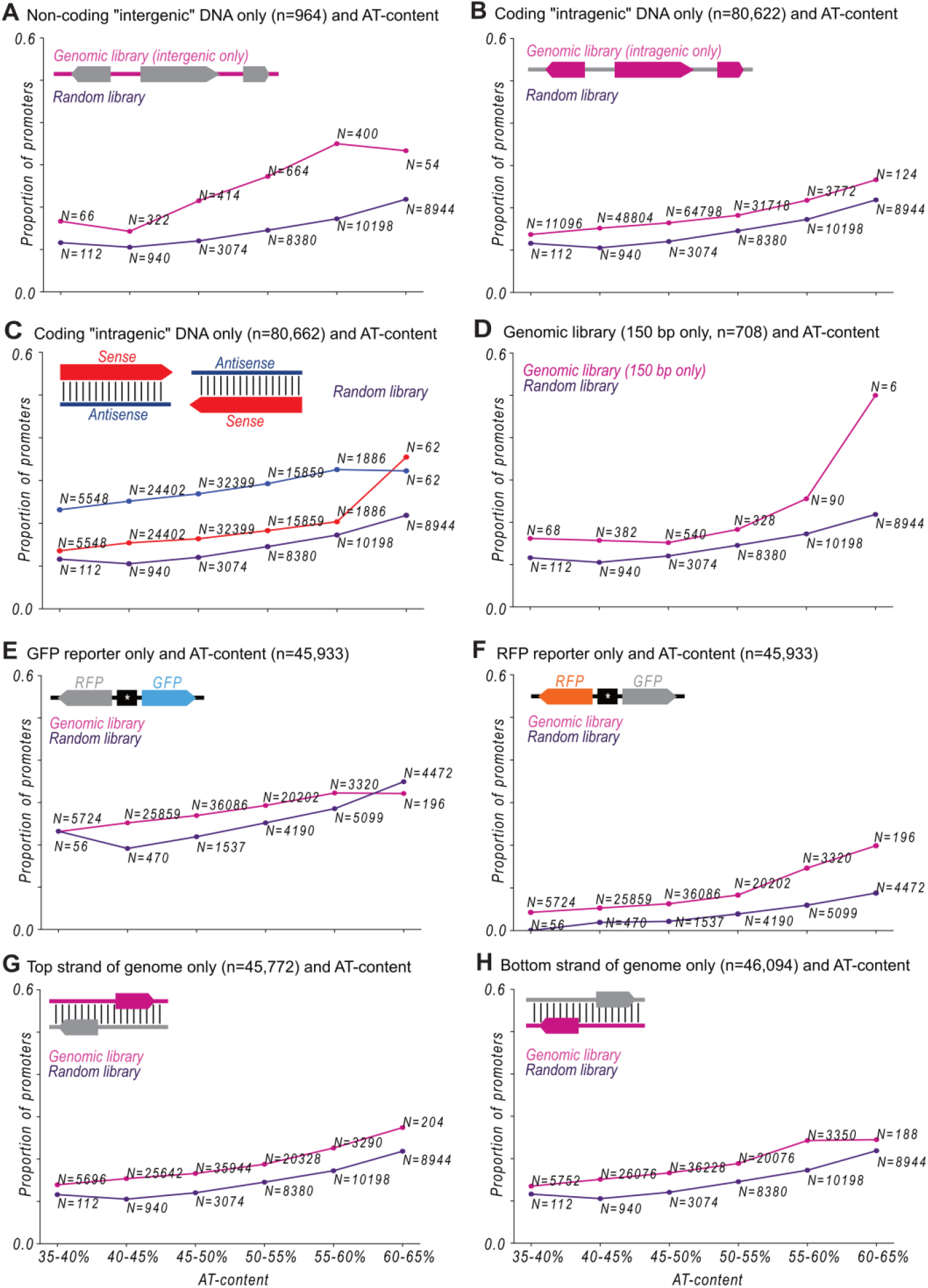
Proportion of promoters and AT-content. (**A**) The proportion of DNA sequences with promoter activity (fluorescence ≥ 1.5 arbitrary units, a.u.) in random DNA (purple) vs in genomic library sequences that lie exclusively in intergenic (non-coding DNA) regions (magenta, n=964 unique sequences). **(B)** Analogous to (A) but comparing random DNA to sequences in the genomic library that lie exclusively in intragenic (inside coding DNA) regions (magenta, n=80,622 unique sequences). **(C)** Analogous to (B) but for the proportion of genomic sequences that lie exclusively inside coding DNA (n=80,622 unique sequences) if they are on the sense-strand (red) or antisense strand (blue). **(D)** Analogous to (A) but comparing random DNA sequences to genomic sequences exclusively 150 bp in length. **(E)** Analogous to (A) but comparing only promoter activity measured from the green fluorescent protein (GFP) fluorescence reporter. **(F)** Analogous to (E) but for the red fluorescent protein (RFP) fluorescence reporter. **(G)** Analogous to (A) but comparing random DNA to genomic sequences that lie on the top strand in their native genomic context. **(H)** Analogous to (G) but for genomic sequences that lie on the bottom strand in their native genomic context. See **Source Data**.

**Fig S3.**
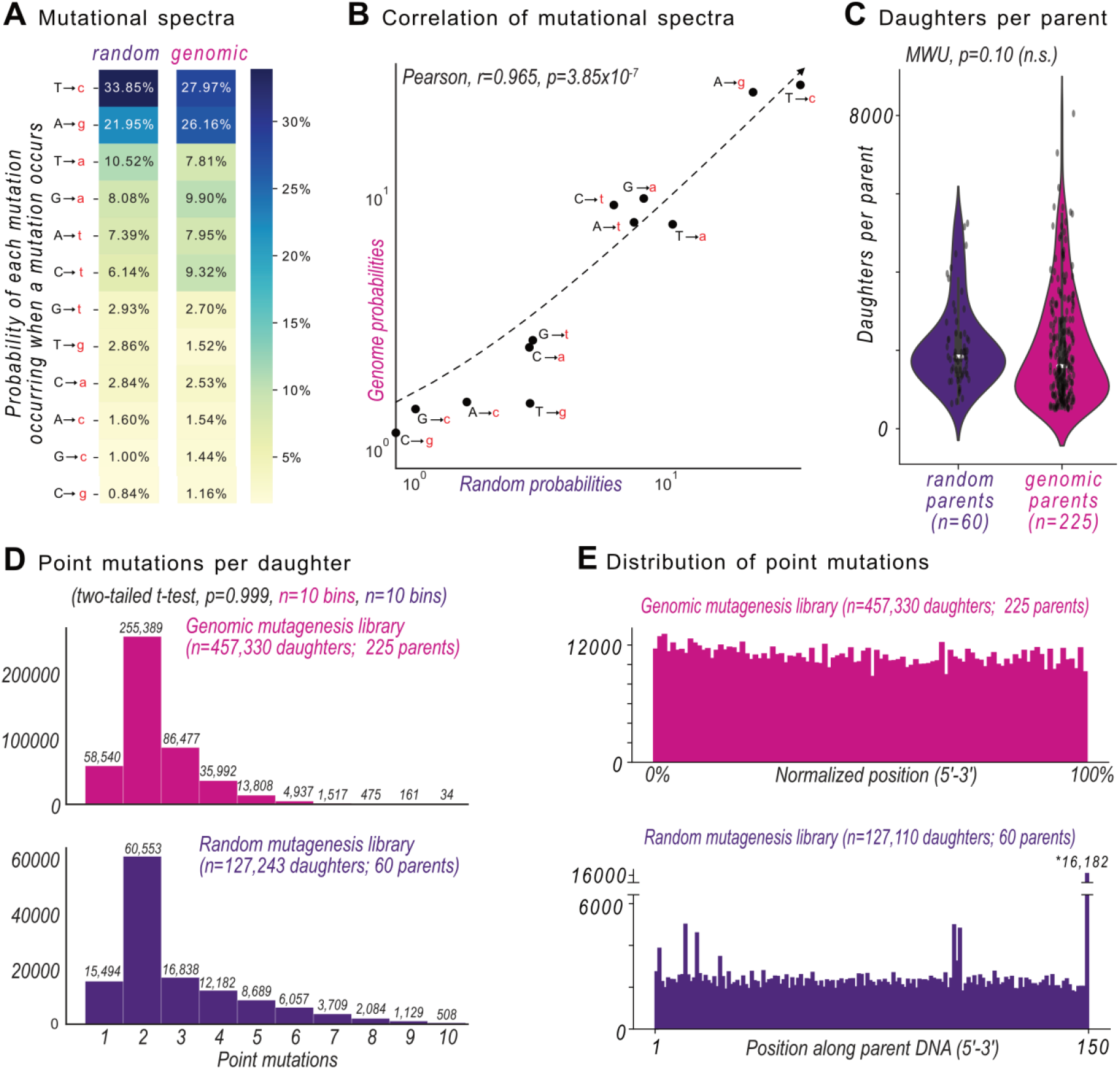
The mutagenesis libraries. **(A)** A heatmap depicting the frequency of different point mutations in the daughter sequences. **(B)** A scatter plot comparing the different point mutation frequencies (see A) in the random mutagenesis library and the genomic mutagenesis library. We tested the null hypothesis that the two frequencies are not correlated (Pearson’s r=0.965, p=3.85×10^−7^). The fitted equation (dashed line) was calculated using the method of least squares. **(C)** The number of daughter sequences for each parent in the random library (purple, left) vs the genomic library (magenta, right; two-tailed Mann Whitney U test, p=0.10, n.s.=not significant). **(D)** Distribution of the number of point mutations in each daughter sequence (top: genome mutagenesis library, bottom: random mutagenesis library). **(E)** The cumulative number of point mutations along the parent sequences, normalized to the percentage of sequence length. Top: genome mutagenesis library, bottom: random mutagenesis library. The reasons why the last position of sequences in the random mutagenesis library experienced more mutations are not known. However, mutual information hotspots do not consistently overlap with the final positions of each parent sequence (see **Fig S5**), suggesting it is not affecting the downstream analyses. See **Source Data**.

**Fig S4.**
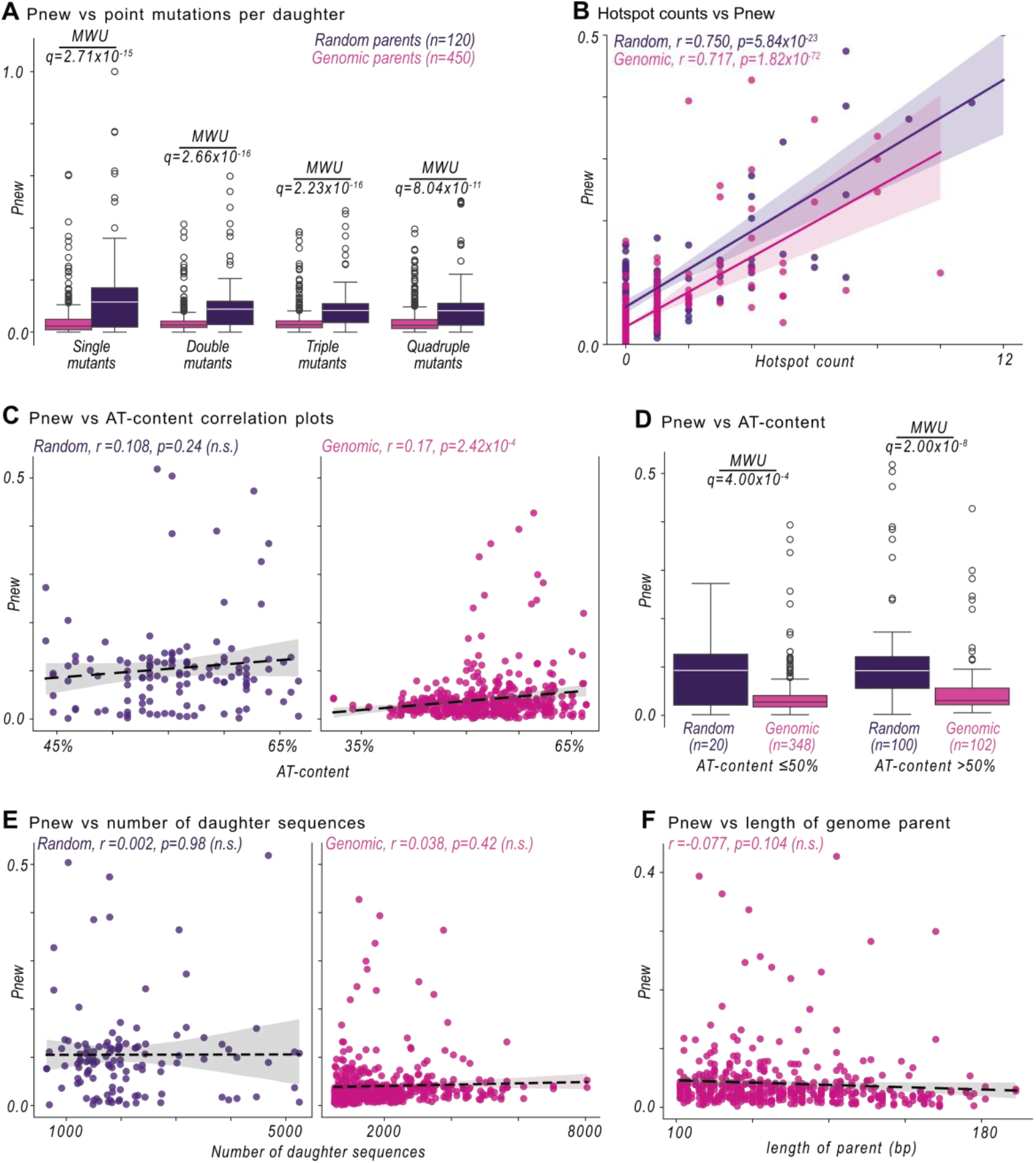
Mutagenesis libraries and Pnew. **(A)** The distribution of Pnew - the probability that a sequence acquires promoter activity by mutation (fluorescence score ≥1.5 a.u., vertical axis) – vs the number of point mutations (horizontal axis) for parent sequences in the genomic mutagenesis library (left member of each pair, magenta) and the random mutagenesis library (right member of each pair, purple). For each box plot, the central line corresponds to the median, the filled rectangle to the interquartile range (IQR), and the whiskers indicate ±1 standard deviation. Outlined circles indicate outliers. For each boxplot pair, we test the null hypothesis that there is no difference in distributions using a two-tailed Mann-Whitney U (MWU) test. We correct the output p-values to q-values using the Benjamini-Hochberg procedure ^42^. Single mutants: q=2.71×10^−15^, double mutants: q=2.66×10^−16^, triple mutants: q=2.23×10^−16^, quadruple mutants: q=8.04×10^−11^. Higher numbers of mutations have too small sample sizes for analysis. **(B)** A scatter plot of the number of mutual information hotspots (vertical axis, see also Fig 2) and Pnew (horizontal axis, see also panel A). The null hypothesis that there is no association between these quantities is rejected (random, purple points: Pearson’s r=0.750, p=5.84×10^−23^; genomic, magenta points: Pearson’s r=0.717, p=1.82×10^−72^). The solid line shows the best-fit linear regression model, with the shaded area indicating the 95% confidence interval. **(C)** Analogous to (B) but for Pnew vs AT-content in the random parents (left, purple, r=0.108, p=0.24, n.s. = not significant) and the genomic parents (right, magenta, r=0.17, p=2.42×10^−4^). **(D)** Analogous to (A) but comparing Pnew in random (purple) and genomic parents (magenta) for those parents with an AT-content ≤ 50% (left pair, q=4.00×10^−4^) and > 50% (right pair, q=2.00×10^−8^). **(E)** Analogous to (C) but for Pnew vs the number of daughter sequences. **(F)** Analogous to (C) but for Pnew vs the length of the parent sequence (bp). See **Source Data**.

**Fig S5.**
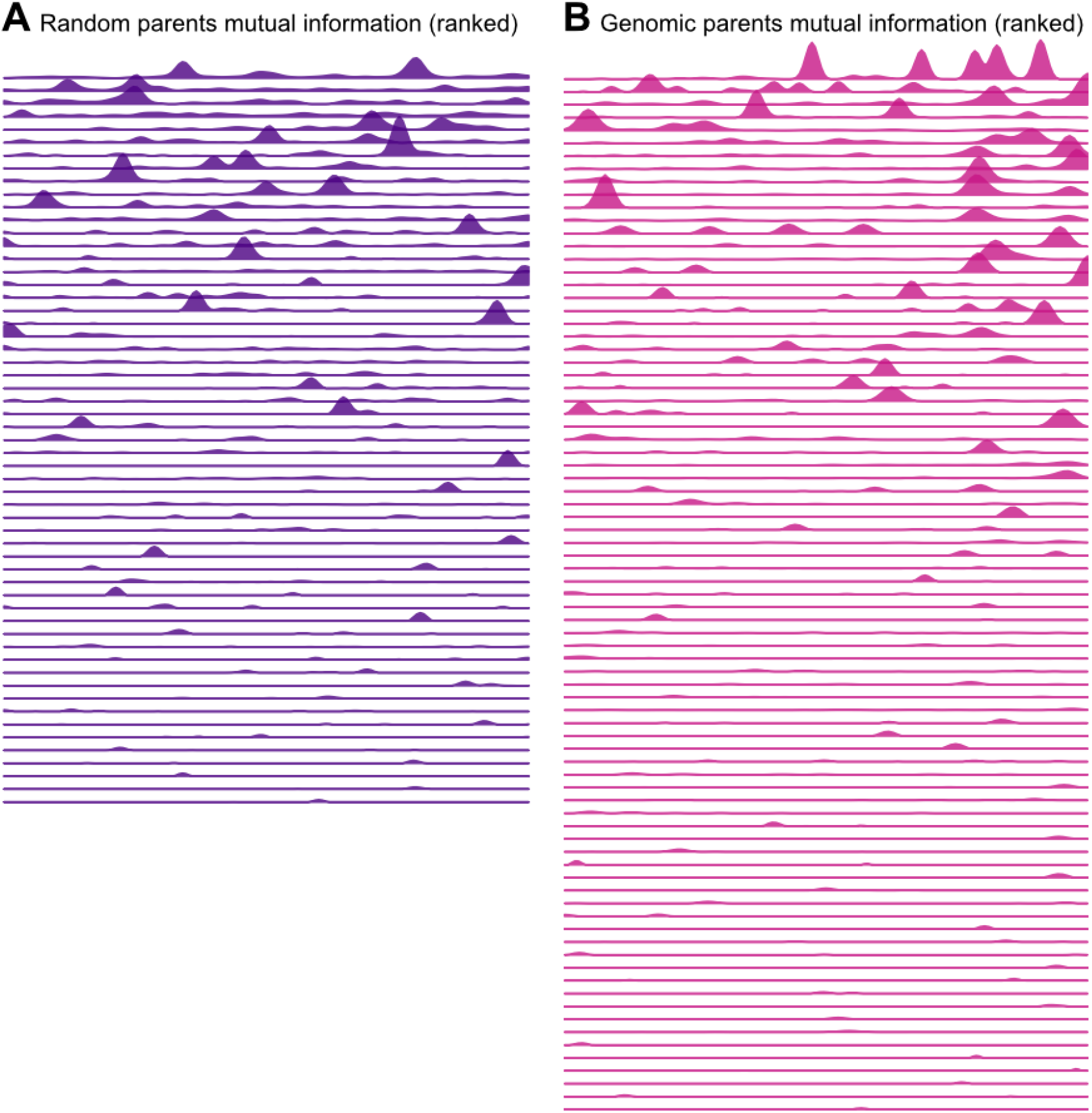
Mutual information hotspots. (**A**,**B**) We calculate at each position (i) along a parent sequence (horizontal axes), the mutual information *I*_*i*_*(b,j)*, vertical axes) between the nucleotide identity *(j* = *A, T, C or G)*, and the fluorescence score *(f* = *1*.*0 – 4*.*0 a*.*u*.) in information theoretical units (bits). Positions are normalized to percentages of parent length, because the genomic parents vary in length. Each plot shows the mutual information of a single parent sequence and fluorophore /genetic orientation. Plots are ranked based on the total sum of the mutual information. Parent sequences without any mutual information hotspots are not shown (see **Methods**). **(A)** Random parents. **(B)** Genomic parents. See **Source Data**.

**Fig S6.**
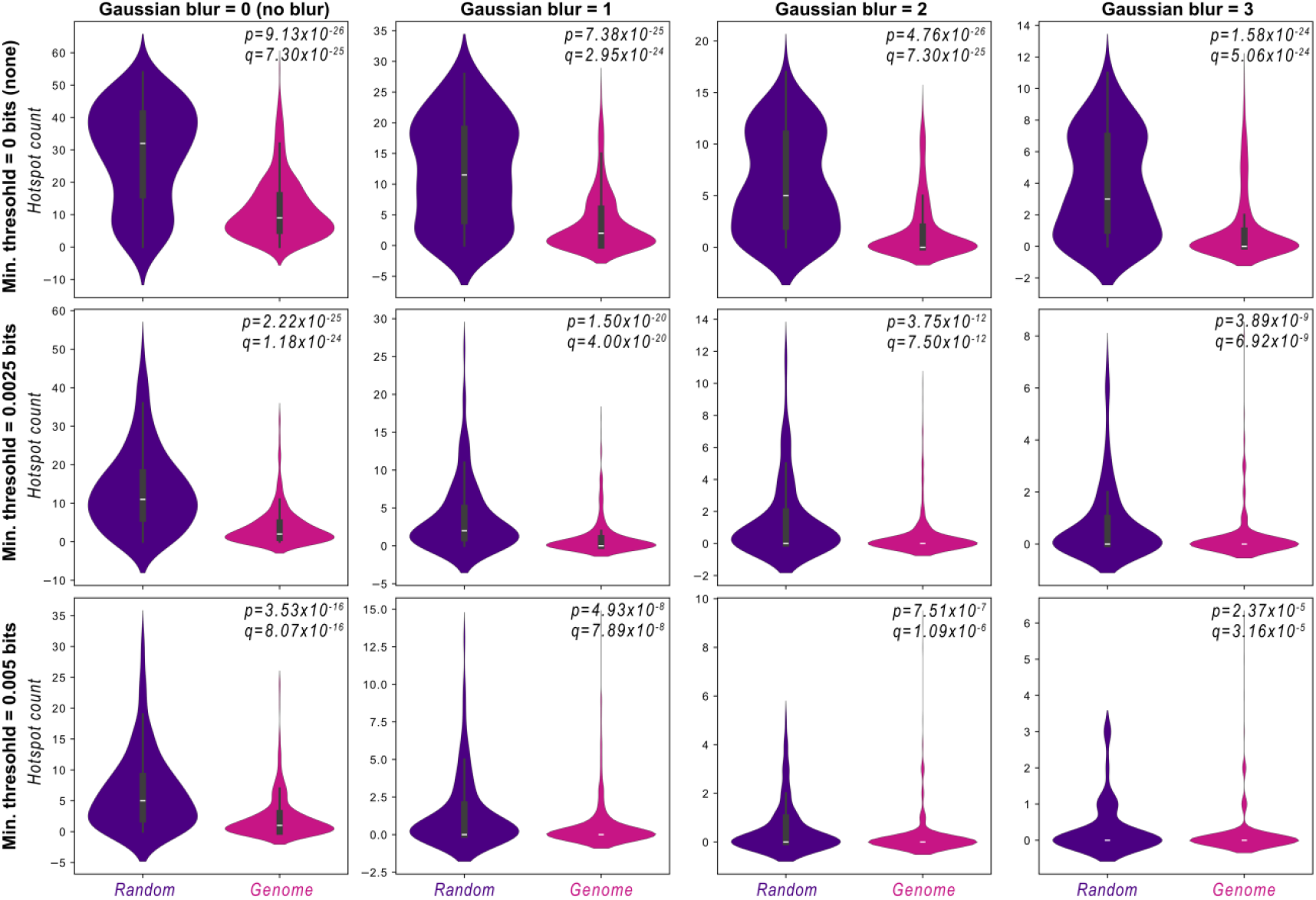
Gaussian blurring and minimal information thresholding vs the number of hotspots. Each panel shows a pair of violin plots that compares the distribution of the number of mutual information hotspots (see also **Fig 2**) between parents in the random library (purple, left plot) and parents in the genome library (magenta, right plot). The white line shows the median, the gray box the interquartile range (IQR), whiskers indicate ±1 standard deviation, and the filled shape reflects a kernel density estimate. For data in each panel, we test the null hypothesis that the distributions are the same, using a two-tailed Mann-Whitney U test to compute a p-value. We account for multiple-hypothesis testing with the Benjamini-Hochberg procedure ^42^ to compute a corresponding q-value. Not significant (n.s.) values are written in red. (Left to right panels) Before calculating the distribution of hotspots, we apply a Gaussian filter (see **Methods**) using the `scipy.ndimage` (v1.13.1) function `gaussian_filter`, incrementally increasing the sigma (σ) parameter for the Gaussian filter from left to right (σ = 0, 1, 2, and 3). (Top to bottom panels) Peaks must be above a minimum threshold to be considered. We incrementally increase this threshold (top to bottom) from 0.000 bits to 0.0025 and 0.005 bits. See **Source Data**.

**Fig S7.**
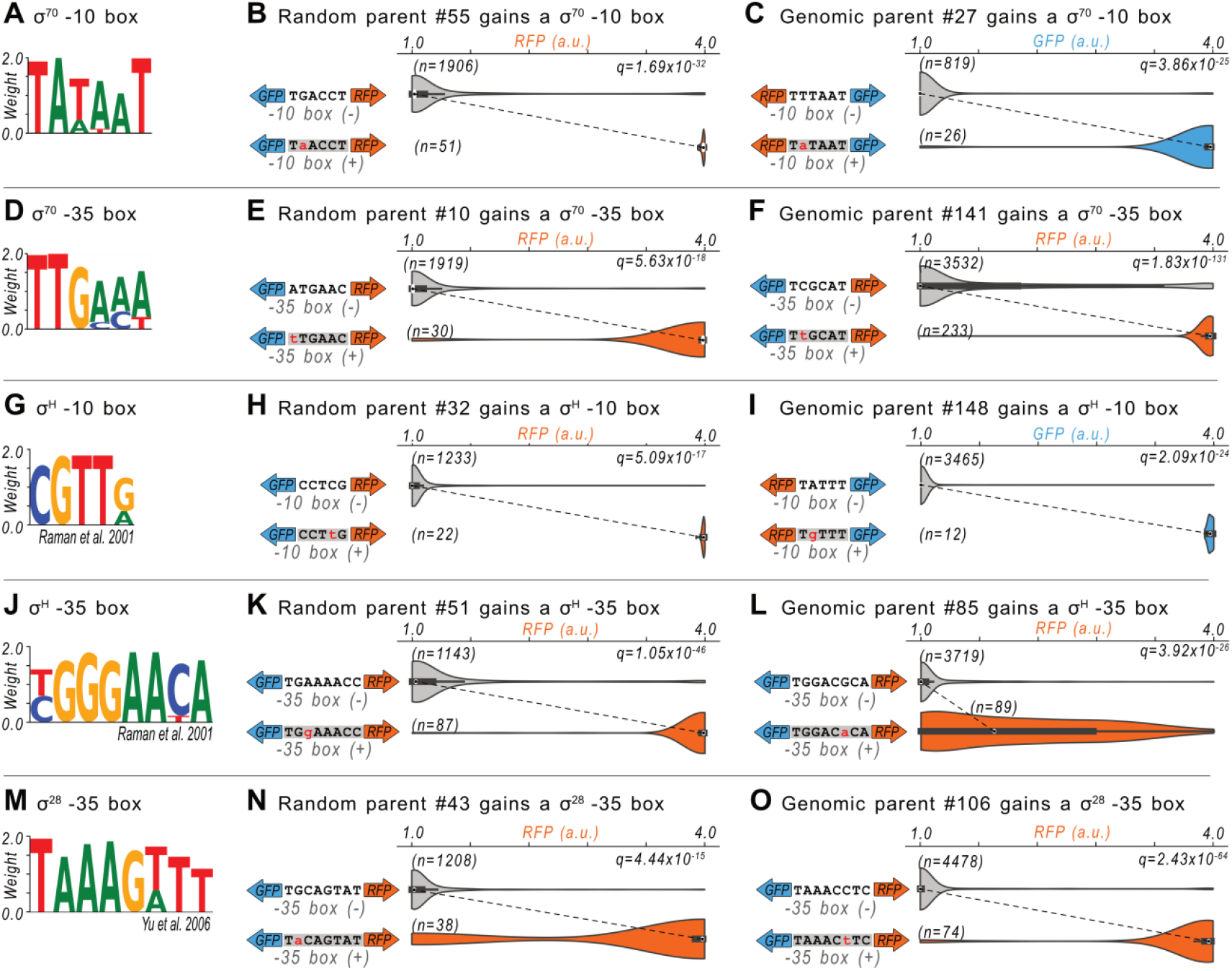
Examples in which mutationally gained sites create promoter activity. **(A)** Sequence logo derived from a position-weight matrix (PWM, see Methods) of a −10 box for the sigma (σ) 70 housekeeping factor. The Logo was drawn by Logomaker^62^. **(B)** Top left: a cropped representation of the DNA of a parent sequence. The arrows indicate the strand and orientation of the downstream reporter gene (teal, green fluorescent protein [GFP], orange, red fluorescent protein [RFP]). Top right: the distribution of fluorescence scores (a.u.) for daughter sequences encoding the sequence at the top left, which is not a predicted −10 box. Bottom left: analogous to the top left. The gray highlighted. DNA sequence corresponds to a gained PWM-predicted site for the transcription factor of interest. Bottom right: analogous to the top right, but for daughter sequences encoding the predicted site of interest at the bottom left. We test the null hypothesis that the two fluorescence distributions are the same with a two-tailed Mann-Whitney U test, and correct the calculated p-values for multiple hypothesis testing using the Benjamini-Hochberg procedure ^42^ with corresponding q-values. Panel (B) specifically corresponds to random parent #55 gaining a σ70 −10 box. **(C)** Analogous to (B) but for genomic #27 gaining a σ70 −10 box. **(D)** Analogous to (A) but for a σ70 −35 box. **(E)** Analogous to (B) but for random parent #10 gaining a σ70 −35 box. **(F)** Analogous to (B) but for genomic parent #141 gaining a σ70 −35 box. **(G)** Analogous to (A) but for a σH −10 box. **(H)** Analogous to (B) but for random parent #32 gaining a σH −10 box. **(I)** Analogous to (B) but for genomic parent #148 gaining a σH −10 box. **(J)** Analogous to (A) but for a σH −35 box. **(K)** Analogous to (B) but for random parent #51 gaining a σH −35 box. **(L)** Analogous to (B) but for genomic parent #85 gaining a σH −35 box. **(M)** Analogous to (A) but for a σ28 −35 box. **(N)** Analogous to (B) but for random parent #43 gaining a σ28 −35 box. **(O)** Analogous to (B) but for genomic parent #106 gaining a σ28 −35 box. See **Source Data**.

**Fig S8.**
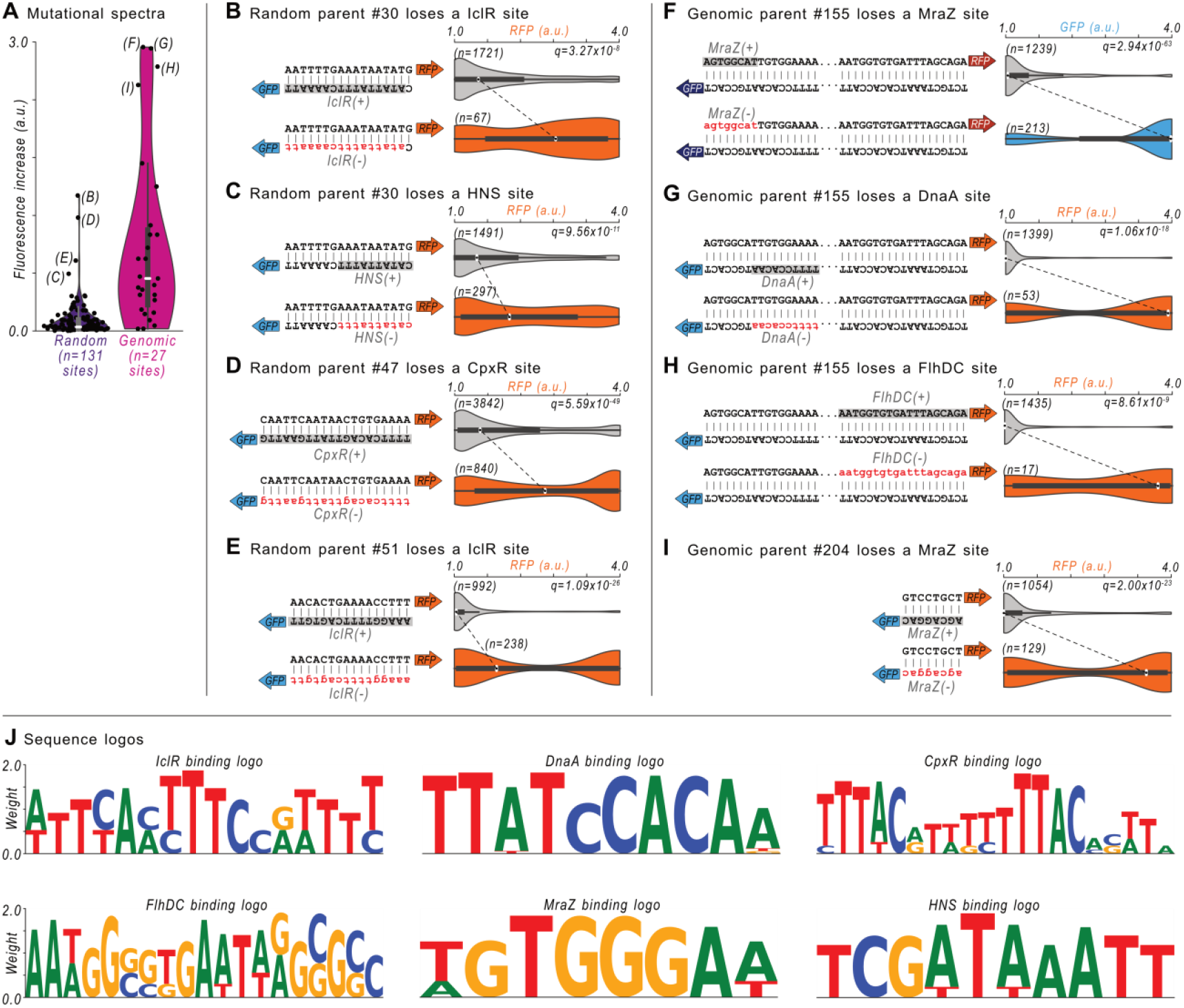
Examples in which losing sites creates promoter activity. **(A)** Fluorescence increases (arbitrary units, a.u.) in parent sequences when a binding site is destroyed, for both random (left, purple) and genomic (right, magenta) parents. The white line shows the median, the gray box the interquartile range (IQR), whiskers span ±1 standard deviation, and the shaded area reflects a kernel density estimate. Black circles correspond to individual data points. Those annotated with letters are the subject of the remaining panels. **(B-I)** Top left: a cropped representation of the double-stranded DNA of a parent sequence. The arrows indicate the strand and orientation of the downstream reporter gene (teal, green fluorescent protein [GFP], orange, red fluorescent protein [RFP]). The DNA sequence highlighted in gray corresponds to a position-weight-matrix (PWM)-predicted binding site for the transcription factor of interest in the wild-type parent. Top right: the distribution of fluorescence scores (a.u.) for all daughter sequences encoding the predicted site of interest. Bottom left: analogous to the top left, but representing DNA sequences no longer encoding the predicted site of interest. Bottom right: analogous to the top right, but for daughter sequences no longer encoding the predicted site of interest. We test the null hypothesis that the fluorescence distributions are the same, using a two-tailed Mann-Whitney U test, and correcting the calculated p-values for multiple hypothesis testing using the Benjamini-Hochberg procedure ^42^ with corresponding q-values. See (A) for details on violin plots. **(B)** Daughter sequences from random parent #30, and their RFP distributions with a IclR(+) and without the IclR(−) site of interest. **(C)** Analogous to B but for losing a HNS site. **(D)** Analogous to (B) but for random parent #47 and losing a CpxR site. **(E)** Analogous to (B) but for random parent #51 and losing a IclR site. **(F)** Analogous to (B) but for genomic parent #155 and losing a MraZ site. **(G)** Analogous to (B) but for genomic parent #155 and losing a DnaA site. **(H)** Analogous to (B) but for genomic parent #155 and losing a FlhDC site. **(I)** Analogous to (B) but for genomic parent #204 and losing a MraZ site. **(J)** Sequence logos derived from the position-weight matrices (PWMs, see Methods) IclR, DnaA, CpxR, FlhDC, MraZ, and HNS from RegulonDB^38^. The Logos were drawn by Logomaker^62^. See **Source Data**.

**Fig S9.**
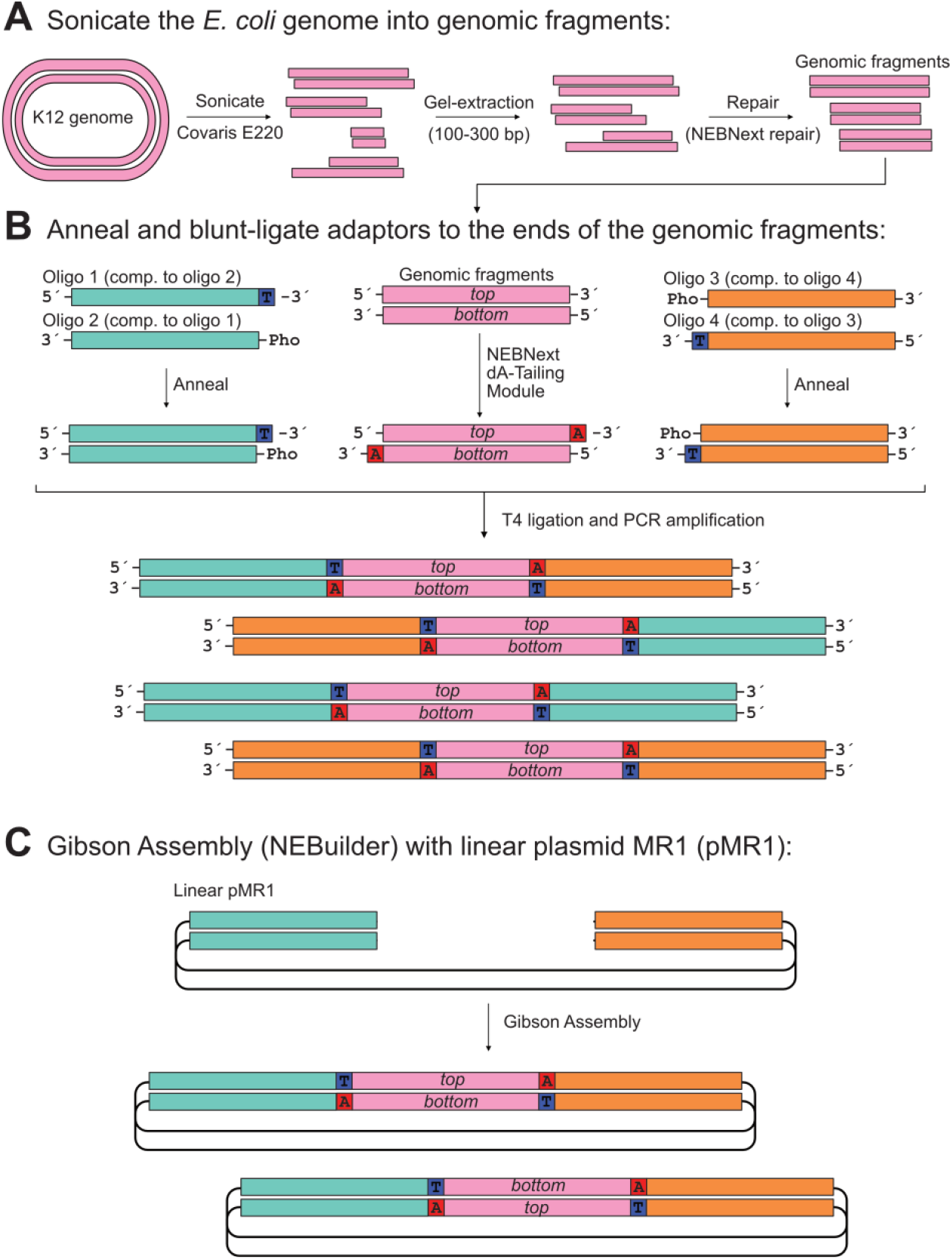
Creating the wild-type genomic library. **(A)** We fragment the *E. coli* genome using an ultrasonicator, isolate sequences of 100-300 bps by gel extraction after electrophoresis, and repair the ends of the fragments using a repair kit. **(B)** Left: complementary (“comp”) oligonucleotides (“oligos”) are annealed. One of the oligos has a thymine (T) overhang on the 3’-end and the other a phosphate (Pho) on the 5’-end. The double-stranded product is homologous to the upstream region of plasmid MR1 (pMR1). Middle: We add an Adenine (A) to the 3’-end of the genomic fragments. Right: analogous to right, except the product is homologous to the downstream region of pMR1. Bottom: double-stranded products annealed using T4 ligation and polymerase chain reaction (PCR). **(C)** A Gibson Assembly reaction with the products from (B) and linearized copies of plasmid MR1 (pMR1) creates circularized plasmid libraries with genomic DNA in either orientation. See **Source Data**.

**Fig S10.**
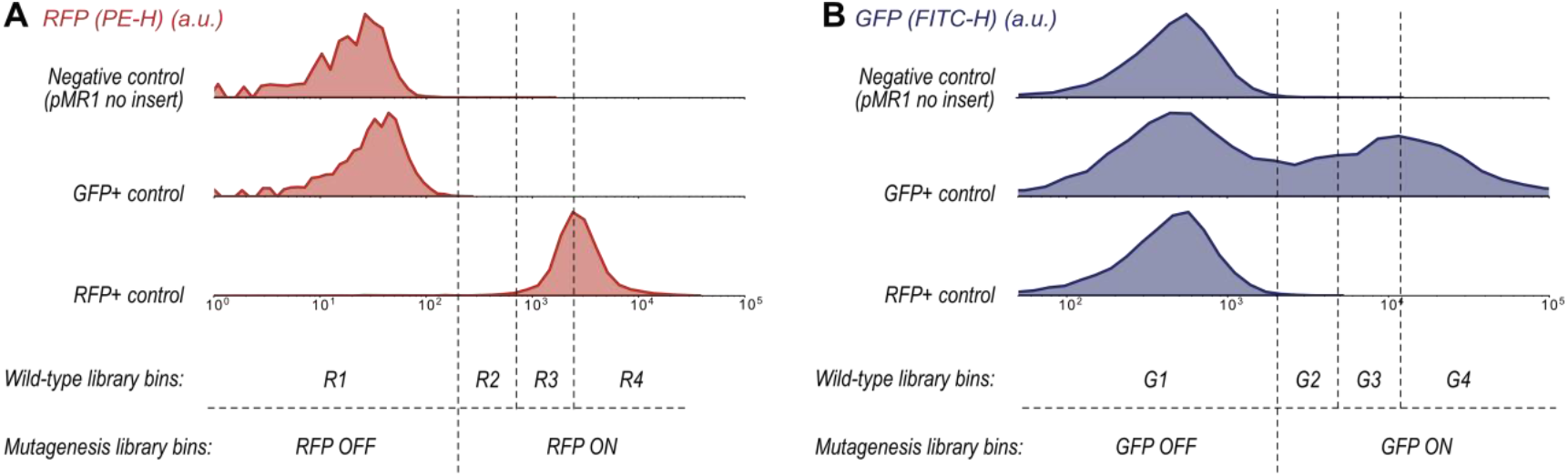
Fluorescence-activated cell sorting boundaries. **(A)** Red fluorescent protein (RFP) readouts in the Phycoerythrin–Height (PE-H) channel for control plasmids. Top: negative control with the plasmid MR1 (pMR1) backbone and no insert. Middle: green fluorescent protein (GFP) positive control consists of the bba_J23110 promoter integrated into pMR1 oriented towards the GFP coding sequence. Bottom: RFP positive control consists of the bba_J23110 promoter integrated into pMR1 oriented towards the RFP coding sequence. The fluorescence distributions determine the fluorescence-activated cell sorting (FACS) bin boundaries, demarcated by vertical dashed lines. The wild-type libraries are sorted into bins R1, R2, R3, and R4. The mutagenesis libraries are sorted into bins RFP-OFF and RFP-ON. **(B)** Analogous to (A) but for Fluorescein Isothiocyanate-Height (FITC-H) and the same control plasmids. The wild-type libraries are sorted into bins G1, G2, G3, and G4. The mutagenesis libraries are sorted into bins GFP-OFF and GFP-ON. See **Methods** and **Source Data**.

